# Track-by-Day: A standardized approach to estrous cycle monitoring in biobehavioral research

**DOI:** 10.1101/2022.08.26.505403

**Authors:** Gianna M. Raimondi, Ashley K. Eng, Murphy P. Kenny, Madison A. Britting, Linnaea E. Ostroff

## Abstract

Despite known sex differences in brain function and incidence of neurological disorders, female subjects are routinely excluded from preclinical neuroscience research, particularly behavioral studies in rats. A common rationale for excluding females is that the hormone fluctuations of the estrous cycle will increase variability in experimental data. Accounting for the estrous cycle as an experimental variable requires expert knowledge of cycle tracking methods, which presents a barrier to widespread inclusion of female subjects. Conventional tracking relies on qualitative interpretation of vaginal cytology smears, and the subjective nature of this approach combined with a lack of reporting standards likely underlies the conflicting literature on estrous cycle effects on behavior. The estrous cycle is traditionally divided into stages based on cytology, but most stages do not directly reflect hormonal events and are therefore of limited relevance to neuroscience experiments. Here we present a simple, streamlined approach to estrous cycle monitoring in rats that eliminates subjective staging. Our method instead indexes the days of the estrous cycle to the one event that is unambiguously reflected in vaginal cytology – the pre-ovulatory surge in 17β-estradiol and subsequent epithelial cornification. With this tracking method, we demonstrate that cycle length is robustly regular across conditions. We quantified long-term memory in a Pavlovian fear conditioning experiment and uterine histology in a large cohort of rats, and found that grouping subjects by day was more sensitive in detecting cycle effects than grouping by traditional cytology staging. We present several datasets demonstrating the logic and applicability of our method, and show that, in the Track-by-Day framework, the cycle is highly regular and predictable in the vast majority of rats across a range of experimental conditions.

## Introduction

Despite well-documented sex differences in many aspects of brain structure and function ^1,2^, female subjects have been largely excluded from most preclinical research in the biobehavioral sciences. Research on rats, a preferred model for behavioral studies, is especially biased towards males; in 2017, 53.5% of studies in high-profile neuroscience journals and 82.2% of studies in behavior-focused journals used only male rats ^3,4^. Anxiety-related disorders are more common in women than men^5,6^, so studying anxiety-related behaviors in female subjects should provide both better insights into brain mechanisms and greater translational relevance. Exclusion of females is often justified on the grounds that hormone fluctuations across the estrous cycle will increase variability in experimental data. Although this concern is controversial ^7–11^, the literature on sex and estrous effects on anxiety-related behaviors is indeed inconsistent ^12–14^. Disagreement between studies can fuel the perception that the estrous cycle is an uncontrollable confound, but it may actually be driven by a lack of universal standards for tracking and reporting the cycle^13^. Conventional methods of tracking the estrous cycle are complex and require expert knowledge^15–17^, so a straightforward, standardized approach tailored for behavioral research is necessary to improve reproducibility between studies and lower a barrier to the use of female subjects.

Sex hormones are known to affect brain function ^18^, and concerns about the estrous cycle in behavioral research relate mainly to two steroid hormones, 17β-estradiol and progesterone, whose serum levels change several-fold over the course of the cycle ^19–21^. Direct measurement of hormones requires blood collection, which is invasive and stressful, so cycle progression is instead monitored by observing changes in vaginal cytology. Traditional tracking methods assign vaginal smears to one of four classic stages of mammalian reproduction: estrus, metestrus, diestrus, and proestrus^22–25^. The stages are defined qualitatively and do not have discrete boundaries, so interpreting smears is an inherently subjective art that requires training and experience ^15–17,26–28^. The rat estrous cycle typically lasts four days, but the four stages vary in length from less than twelve hours to more than two days. Nevertheless, the stage names are sometimes used to refer to days of the cycle and studies often do not specify which is meant, resulting in inconsistent and broad temporal ranges between studies for a given stage. It is standard to exclude subjects with irregular cycles from experiments, but there is no standard definition of regular cycling. Exclusion criteria are therefore subjective, and perceptions that the cycle is highly sensitive to environmental conditions^16^ and that irregularity is common^15^ discourage estrous cycle tracking altogether. Semi-quantitative approaches ^29–31^ and machine learning tools ^32,33^ can reduce subjectivity in cytology interpretation, but a rigorous reporting framework and simpler methods that are accessible to the uninitiated are still needed.

The implicit rationale for using cytology stage as a grouping factor in behavioral research is that the stages reflect serum levels of neurobiologically-relevant hormones. The stages were defined long before the hormone cycle was understood, however, and only one change in the vaginal epithelium has been demonstrated to be directly and temporally linked to a hormonal change: desquamation of the stratum corneum the day after the preovulatory 17β-estradiol surge ^34,35^. This event defines one of the four cytology stages: estrus. Since there is no evidence that the other stages carry experimentally meaningful information, a more effective and streamlined approach would be to define the cycle by days relative to the known peri-ovulatory cytology. To compare this Track-by-Day method to traditional staging, we quantified cytology across cycles under several conditions. Cycle length was robustly regular and predictable when indexed by day, whereas the progression of cytology stages was highly variable. Interestingly, we found effects of the estrous cycle on Pavlovian fear conditioning that were specific to either cycle day or traditional cytology stage. We also found that cycle days are better correlated with hormone-sensitive features of uterine histology than stages. The Track-by-Day approach is simple, highly accessible, and can be used in conjunction with, or instead of, traditional stage assignments to detect effects in females that may be otherwise obscured.

## Results

### An effective strategy for the use of vaginal cytology as a proxy for hormone cycles

Levels of circulating sex hormones across the rat estrous cycle were quantified by several studies in the 1970’s ^19–21^. These studies found strikingly similar patterns for all hormones, including 17β-estradiol and progesterone (Figure 1a). Both hormones are at their lowest on the day of ovulation, and 17β-estradiol begins a gradual rise on the third day, then surges to a peak on the fourth day. In four-day cycles, 17β-estradiol falls rapidly on the fourth day and a sharp peak in progesterone that afternoon precedes ovulation. Five-day cycles are less common and differ in that the 17β-estradiol peak is extended by a day and the ensuing progesterone peak is higher ^19^. Since the hormone cycle is stereotyped, it should be possible to extrapolate its timing if a single event can be unambiguously detected. The effects of 17β-estradiol and progesterone on the vaginal epithelium (Figure 1b) confirm that vaginal cytology should be useful for this purpose. 17β-estradiol and progesterone induce terminal squamous and mucosal differentiation, respectively, with 17β-estradiol acting on a faster timescale ^34^. The preovulatory surge in 17β-estradiol induces epithelial cells to proliferate and keratinize, creating a cornified cell layer (stratum corneum), accompanied by leukocyte infiltration into the upper layers of the epithelium. Keratinization ceases with the fall in 17β-estradiol, the progesterone peak induces maturation of mucosal cells, pushing the stratum corneum towards the surface to be delaminated or shed, and the epithelium then enters an atrophic state until 17β-estradiol rises again ^25,34,36–38^.

**Figure 1.**
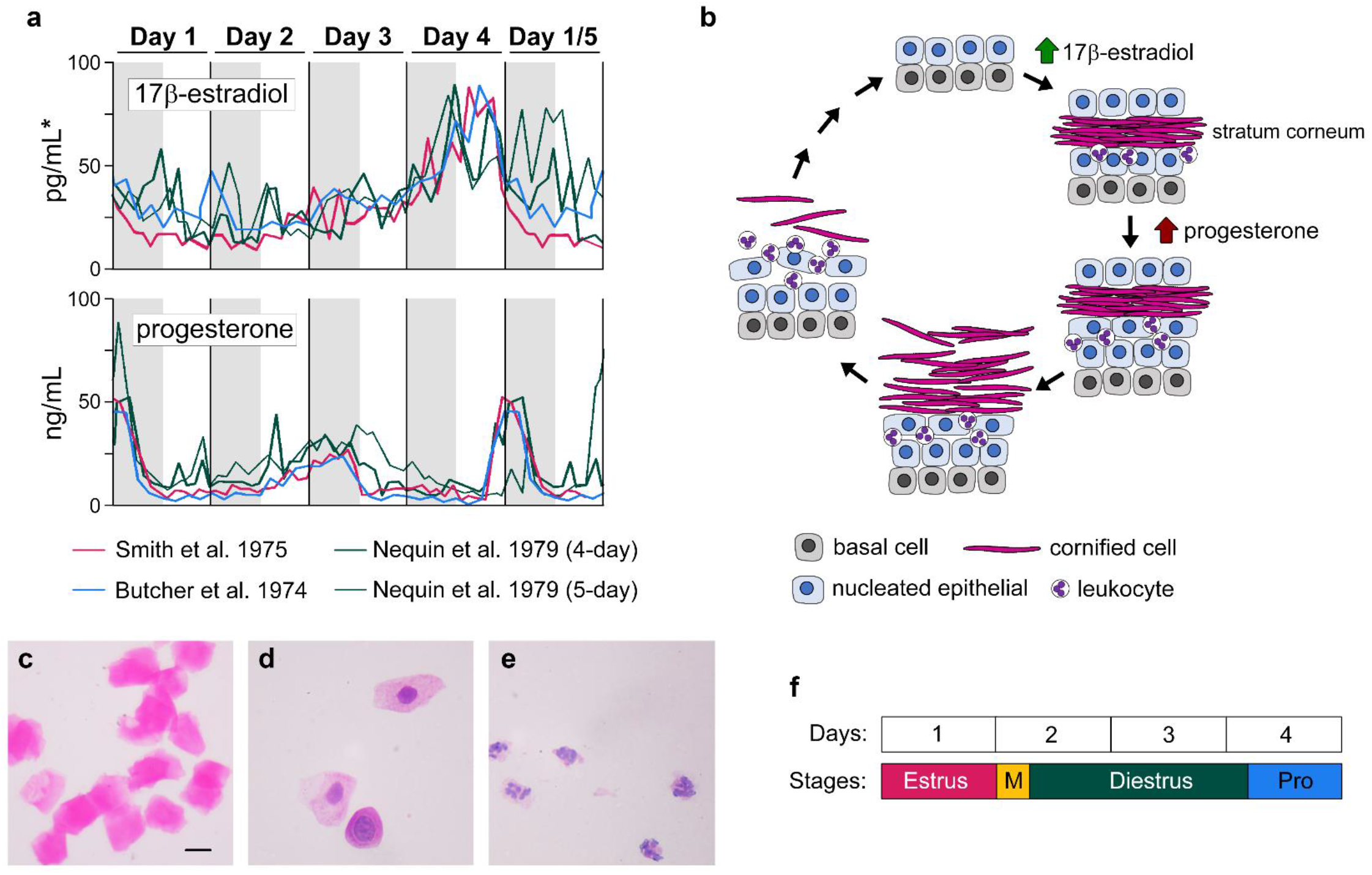
Estrous cycle progression. (**a**) Serum concentrations of 17β-estradiol and progesterone over four- and five-day cycles in the rat as reported in three studies. Data adapted from Butcher et al. (1974), Smith et al. (1975), and Nequin et al. (1979). Gray regions indicate the 12-hour dark period of each day, except for the Nequin et al. study which used a 10-hour dark period. Day 1 is the day of ovulation and termed the estrus day in all studies. *Axis values from Ref. 21, with data from Refs. 19 and 20 plotted at 66% and 200% respectively. (**b**) Schematic of changes in the vaginal epithelium in response to the brief surges of serum hormones during the estrous cycle. (**c-e**) Examples of the three cell types in a vaginal smear stained with H&E: (c) cornified, (d) nucleated epithelial, and (e) leukocytes. Scale: 10µm. (**f**) The four terms that denote the progression of the rat estrous cycle may be used to refer to days (top row) or to cytology stages (bottom row).

Traditional cytology staging takes the entire cycle of the vaginal epithelium into account by evaluating the relative representation of cell types in daily smears ^16,17,26–28,31^. Three general types are seen: cornified cells (Figure 1c), which are flattened, enucleated, and fully keratinized; nucleated (non-keratinized) epithelial cells (Figure 1d); and leukocytes (Figure 1e), which are small and have multi-lobulated nuclei. Smears dominated by cornified cells, reflecting delamination of the stratum corneum, define the estrus stage. Estrus cytology is seen on the day of ovulation, or the day after the peak in 17β-estradiol ^19–21^. Once the stratum corneum has sloughed off, lingering leukocytes and nucleated epithelial cells appear in smears. Prevalent leukocytes define the metestrus stage, and their disappearance relative to nucleated epithelial cells marks the diestrus and proestrus stages, which differ more subtly. Estimates of stage duration vary broadly, but metestrus is generally agreed to be the shortest and diestrus the longest (Figure 1f). Some authors omit metestrus as a separate stage, and others divide diestrus into diestrus I and diestrus II^13,16^.

The timing of the stage transitions can create spurious comparisons if they are used as categorical factors. There is no hormonal event that justifies dividing the second cycle day into metestrus and diestrus, and the hormone peaks on the fourth day are not reflected in a change from diestrus to proestrus cytology, but instead in estrus cytology on the following day. Conversely, diestrus encompasses the second day, when hormone levels are low, through at least part of the fourth day, when they are high. Desquamation of the stratum corneum, indicating the stage of estrus, is often considered to last roughly one day. A simpler and more meaningful strategy would be to use cornified cell desquamation to detect the first cycle day, then account for the cycle temporally instead of ambiguous cytology stages. To determine whether this approach would be viable, we tested its fundamental requirements: unambiguous, temporally discrete cytology and predictable cycle duration.

### Cornified cell desquamation is easily detected in vaginal smears

The desquamation of the 17β-estradiol-induced stratum corneum must be unmistakable if it is to be used to time the cycle. Histological stains provide contrast between cell types on vaginal smears and are often used to facilitate estrous staging ^26,30,39^, so we evaluated three stains – hematoxylin and eosin (H&E), Shorr stain, and cresyl violet – for their ability to distinguish cornified cells from other types. Even without staining, smears containing clumps of delaminated cornified cells (Figure 2a) could be distinguished from smears containing a more disperse mix of cell types (Figure 2e). H&E staining allowed rapid, unambiguous differentiation of smears consisting entirely of cornified cells (Figure 2b) from smears containing a mix of cell types (Figure 2f) due to the contrast between the bright pink cornified cells (Figure 1c) and the purple nuclear staining of the other cells (Figure 1d-e). Shorr stain also readily revealed contrast between cornified cells (Figure 2c) and other cell types (Figure 2g). Although cornified cell clumps were visible with cresyl violet staining (Figure 2d), it did not produce chromatic contrast between cell types (Figure 2h), and thus offered only a marginal improvement over unstained smears. We chose to stain with H&E for the majority of experiments, since it produces clear contrast and is more widely used in laboratories than Shorr stain.

**Figure 2.**
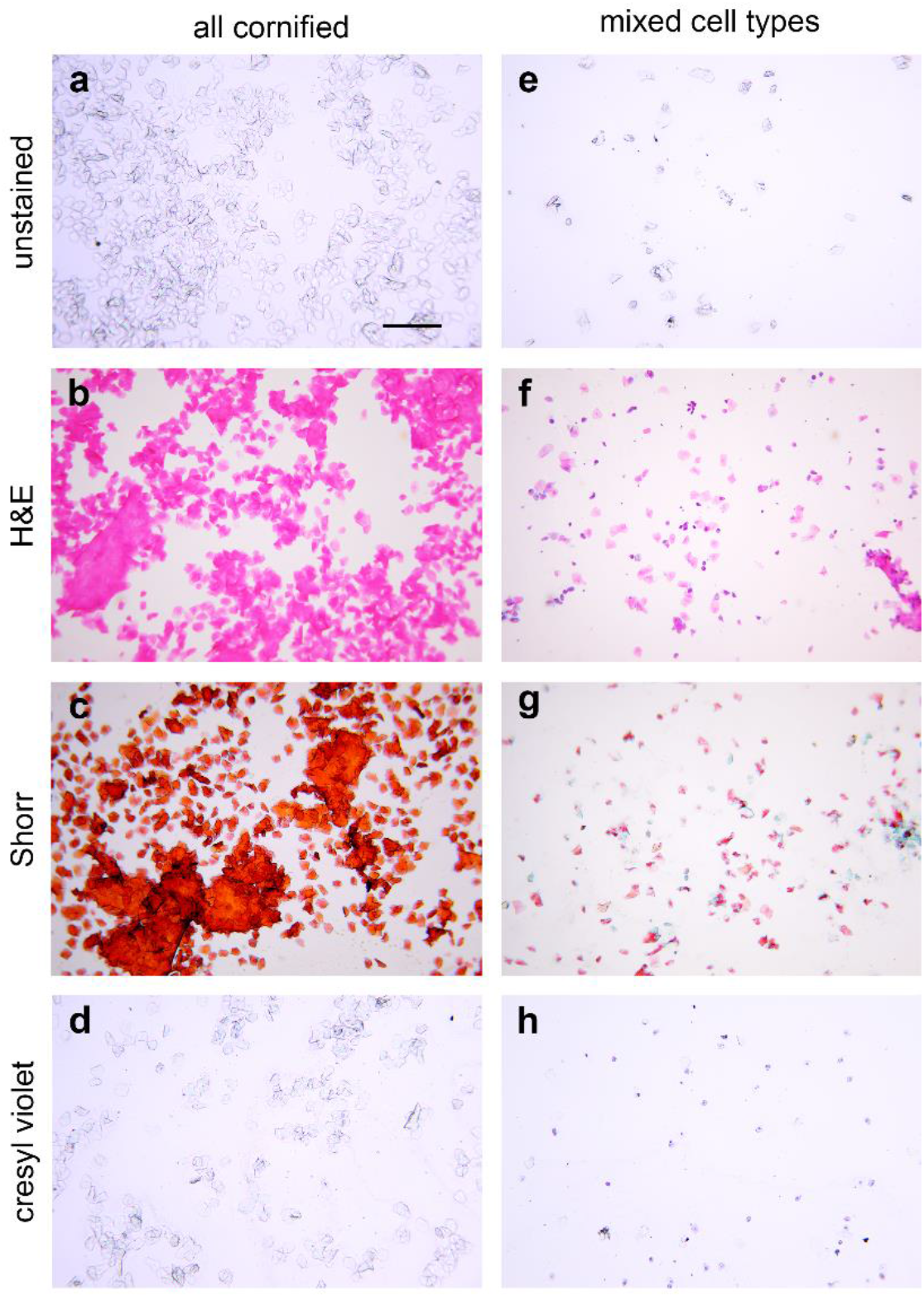
Vaginal cytology under different staining conditions. (a-d) Smears consisting only of cornified cells. (e-h) Smears containing a mix of cell types. Contrast between cell types is low in unstained smears (a,e), high in smears stained with hematoxylin & eosin (H & E; b,f) or Shorr stain (c,g), and low in smears stained with cresyl violet (d,h). Scale: 100µm.

### Full cornification of vaginal smears peaks regularly and universally across subjects and cycles

To determine whether cornified cell desquamation could be used to time the cycle, we collected vaginal smears by swabbing twelve adult (PND 92) female rats for eight days. Animals were housed in ventilated cages in a breeding colony with a 14-hour light period, and smears were collected once daily between one and two hours after lights-on. The percentage of each cell type was quantified for each smear. Cornified cells were present in every smear and in some were the only cell type present, consistent with bulk desquamation of the stratum corneum (Figure 2a-d). Within the first three days, a single read of 100% cornified cells (cornified peak) was seen in all twelve rats (Figure 3a). The cornified peak did not occur on the same calendar day for all animals, but when reads were aligned by the first cornified peak, a second peak occurred four days later in eleven of the twelve subjects (Figure 3b). Cornified peaks occurred simultaneously in only two of the six cage-mate pairs, consistent with reports that rats do not synchronize cycles^40^. We designated the day of a cornified peak as Day 1 and numbered the other days of the cycle accordingly. Eight consecutive days of swabbing ensured each cycle day occurred twice. When the percentage of cornified cells on each read was averaged by cycle day, it was higher on Day 4 than on Days 2 and 3, but variability was high on all three days (Figure 3c). To examine within-rat variability across cycles, the absolute value of the difference between the two cycles was calculated for each cycle day. Day 1 was the most consistent between cycles, as expected, and Day 4 the least (Figure 3d).

**Figure 3.**
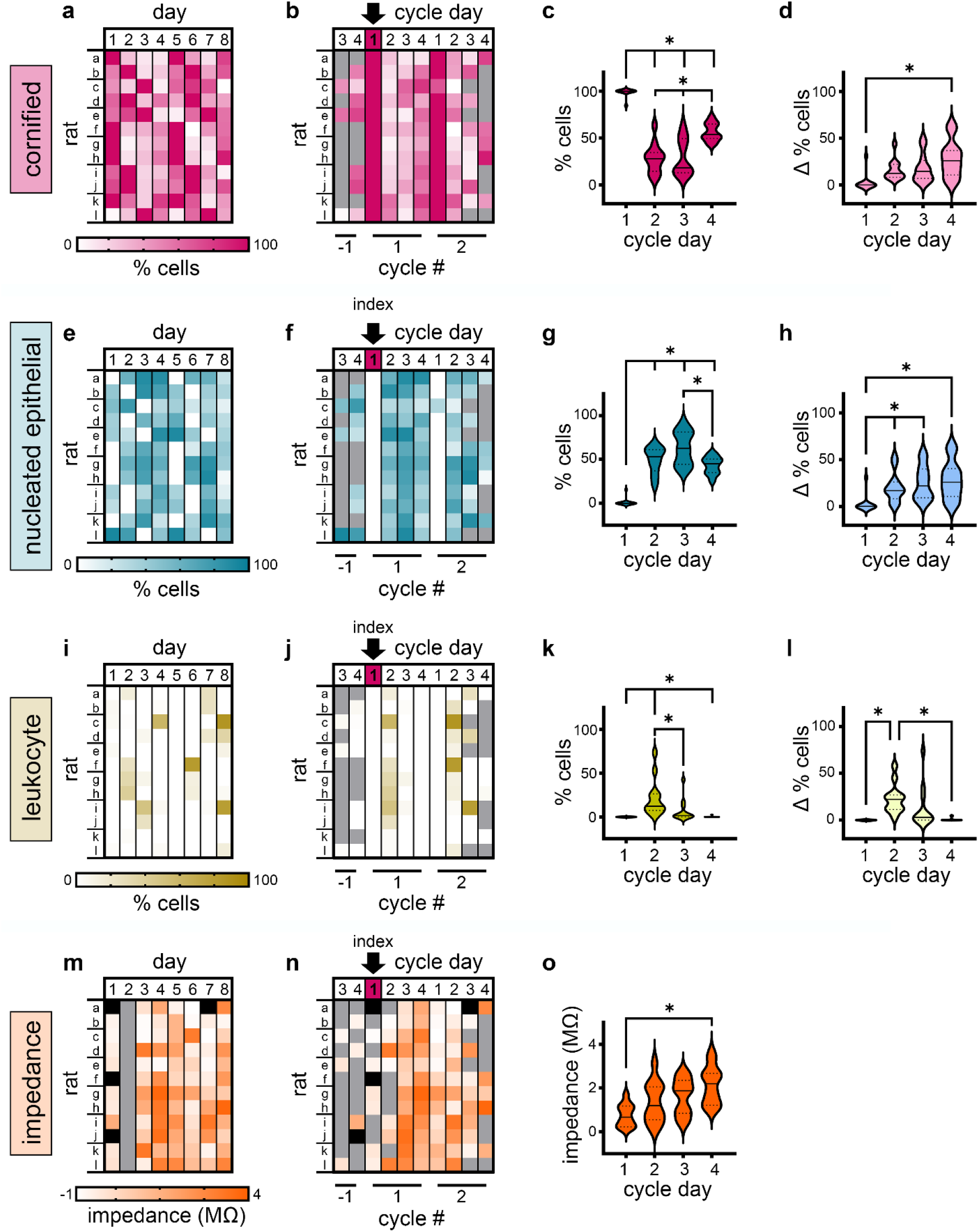
Vaginal cell types across days. (a) Cornified cells as a percentage of all cells for twelve rats over eight days. Gradient scale also applies to (b). (b) Progression of cornified cell percentage aligned by each rat’s first peak (100%). (c) Percent cornified cells by cycle day, averaged averaged across rats according to cycle day. (d) Absolute value of the difference in % cornified cells between the two cycles. (e-l) Data for nucleated epithelials (e-h) and leukocytes (i-l) in the same format as data in (a-f). (m) Vaginal impedance (MΩ) collected on the same eight days. Impedance readings were not taken for the second day of tracking (gray), and a few individual readings were inconclusive (black). Gradient scale also applies to (n). (n) Impedance measures aligned by the day of the first cornified peak as in (b). (o) Mean impedance on each day of the second cycle only. For all raster plots, lines on y-axis indicate cage-mate pairs. * p<0.05

Nucleated epithelial cells were present in most smears on Days 2-4 (Figure 3e) but showed high variability when aligned by the cornified peak (Figure 3f) with the highest percentage on Day 3 (Figure 3g). Within-rat variability was similarly high after Day 1 (Figure 3h). Leukocytes were detected less often than the other cell types (Figure 3i). Some cycles did have a surge in leukocytes on Day 2 (Figure 3j), consistent with the definition of metestrus, but this did not occur in all rats (Figure 3k) and was not consistent within rats across the two cycles (Figure 3l). Vaginal impedance has been reported to change over the estrous cycle and has been advocated as an easy alternative to cytology for cycle tracking^41–44^, although one study found no correlation between impedance and serum hormone levels ^45^. We measured impedance at the time of most swab collections and found that it varied over the cycle in all animals (Figure 3m). No clear cyclic pattern relative to the cornified cell peak was observed (Figure 3n), although on average impedance was lower on Day 1 than Day 4 (Figure 3o). Within-subject comparison of the two tracked cycles shows that Day 1 cytology is overwhelmingly the most predictable aspect of the cycle (Figure 3p).

### The cornified cell peak is regular across environmental conditions

The animals in our first experiment were housed with a 14-hour light period, which is common in breeding colonies but not typically used for behavior experiments. To ensure that the cornified peak is reliable in animals housed on a 12-hour light cycle, we began with a new cohort of 16 rats (PND 35). These animals were housed in non-ventilated cages in a room with only female rats and nursing litters. To determine whether cytology would be more predictable early or late in the day, swabs were collected twice per day for eight days, at two hours and eight hours after lights-on (L2 and L8). For the next eight days, swabs continued to be taken from 11 of the 16 rats. Impedance was measured and blood samples were collected from the lateral tail vein alongside many of the swabs in the second round. For the last four days, swabs were collected from all 16 rats once per day, six hours into the light period (L6). The proportion of each cell type was quantified on every smear (Supplemental Figure 4-1). During twice-daily collection, all 16 rats had regular, discrete cornified peaks consisting of 1 – 4 consecutive reads. Most four-day cycles had either one peak read at the L2 time point or two peak reads, usually L2 followed by L8. This pattern indicates that the morning (L2) read is more reliable than the afternoon (L8) read for tracking the cycle, and suggests that the cornified peak lasts about 24 hours and begins sometime during the dark period. Because some rats had a cornified peak on the first day of sampling, we aligned the cornified cell data to the second peak to better visualize the distribution of surrounding reads (Figure 4a). No patterns were evident in the other two cell types or in impedance (Supplemental Figure 4-2), so only the cornified cells were used to track cycles. The intent of the blood collection was to verify levels of 17β-estradiol and progesterone, but tail vein sampling did not yield sufficient volumes of serum for either of two commercial ELISA kits we tested.

**Figure 4.**
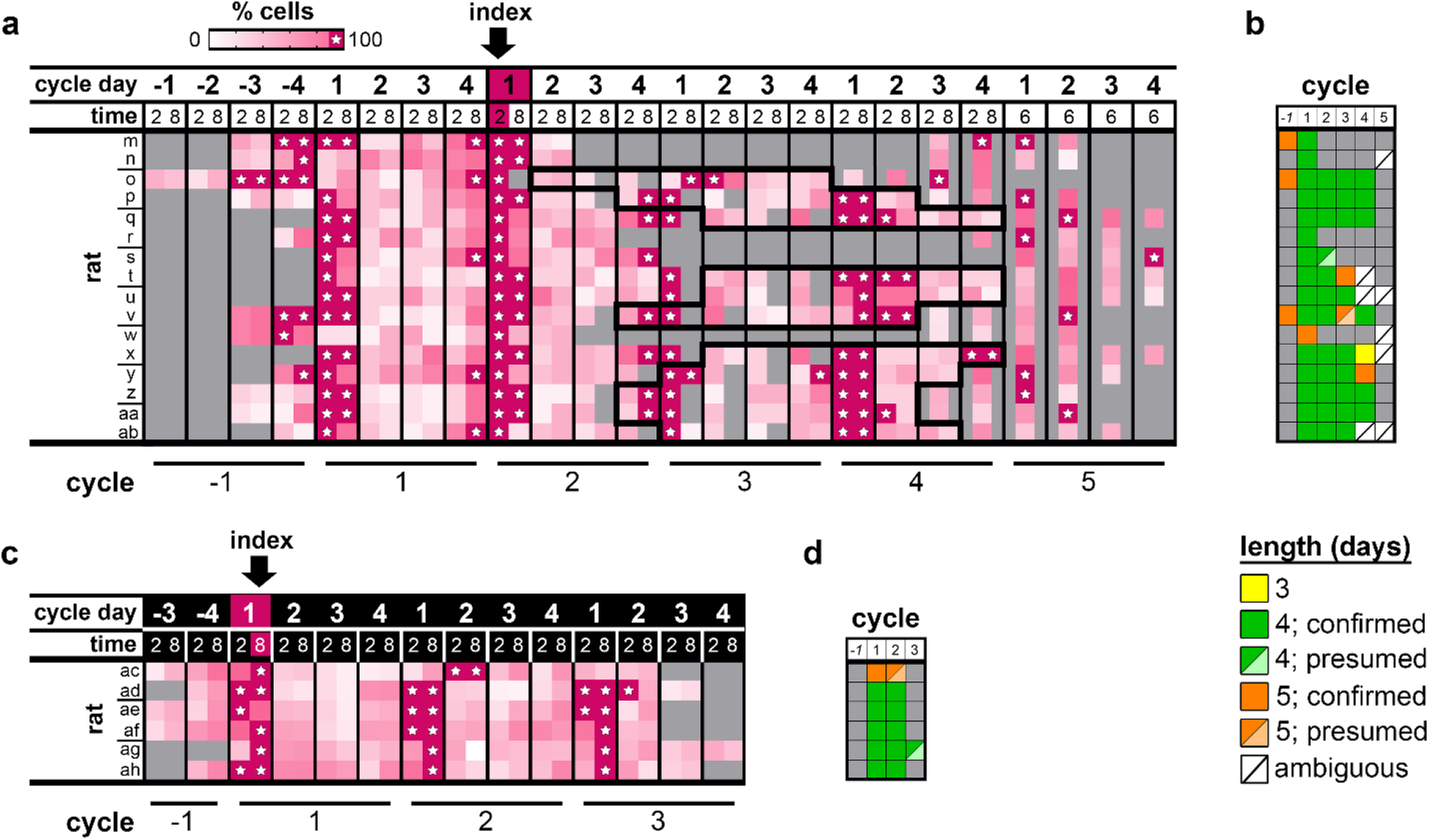
Tracking cornified cells across the light-dark cycle. (a) Percent cornified cells in smears collected at time points 2, 6, or 8 hours after lights-on from rats housed on a 12-hour light-dark cycle. Reads are indexed to the lights-on + 2h peak read of each rat’s second tracked cycle. Times when a smear was not collected are shown in gray. The black outline denotes days when tail vein blood draws were also performed on 11 of the 16 rats. (b) The majority of tracked cycles lasted four days, and every rat had at least one four-day cycle. A confirmed 4-day cycle was one in which all four days plus Days 1 and 2 of the next cycle were observed. A 4-day cycle was presumed if the following Day 2 smear was not available to exclude the possibility of a 5-day cycle. 4-day cycles were also presumed when the first or final few days of tracking were consistent with a partial 4-day cycle. (c) Percent cornified cells in smears collected twice per day, 2 and 8 hours after lights-off, and indexed to the first peak 8-hour read. (d) Five of six animals had two consecutive four-day cycles, and the sixth had a five-day cycle followed by a second apparent five-day cycle. Lines on y-axis indicate cage-mate pairs.

Most cycles lasted four days, but in contrast to the first experiment some five-day cycles were observed. Five-day cycles are generally characterized by two consecutive days of cornified peaks, consistent with the extended peak of 17β-estradiol reported for five-day cycles^19^. Clear five-day cycles were observed in five of the 16 rats, but all five also had four-day cycles (Figure 4b). Four of the eleven rats that underwent blood draws shifted their cycle length by one day, and some did not have detectable peaks in the final round of smears that were collected at the L6 time point. This could reflect disrupted cycles, but it is also possible that peaks were missed if their timing was shifted away from mid-day. Peaks occurred on the same days for two of the eight cage-mate pairs, again demonstrating chance levels of synchrony within cages.

### The cornified peak is reliably detected during the dark period

Rats are nocturnal, so behavior experiments performed in the dark period may be more ethologically relevant. To confirm that the cornified peak could be used to time the cycle during the dark period, we housed a new cohort of six rats (PND 41) in a room with a reverse 12-hour light cycle. This room contained a breeding colony and had ventilated cages, as in the first experiment. Smears were collected twice per day at two hours and eight hours after lights-off (D2 and D8) for twelve days, and cells were quantified for each smear (Supplemental Figure 4-3). Cornified peaks were detected more often in the D8 smears than in the D2 smears, so cycles were aligned to the first D8 peak read (Figure 4c). When two consecutive peaks occurred, the first was more often a D2 read. This order plus the higher frequency of peak reads at D8 and L2 versus D2 and L8 indicates that the cornified peak commences around the light-to-dark transition. Only one of the six rats had a five-day cycle, versus eight four-day cycles amongst the other rats (Figure 4d). Interestingly, two-day peaks were not observed in this group. Leukocytes were also very scarce relative to the smears collected during the light period, consistent with the reported short duration of the metestrus stage, which is defined by copious leukocytes.

### Track-by-Day is more sensitive than traditional staging in detecting estrous effects on fear conditioning

We next compared the performance of the Track-by-Day method to traditional staging in a Pavlovian conditioning paradigm to compare fear learning across days of the estrous cycle, and between females and males. Smears were collected from females once per day, mid-dark period, for two rounds of 8-12 days, with a break of 8 or 10 days in between. Multiple tracking rounds allowed each subject’s cycle to be observed over four weeks, and the break allowed us to interleave 60 female rats in a large experiment. Each smear was classified as either 100% cornified, signifying Day 1, or not. In a few cases, a read of 90-99% cornified cells was used to define Day 1 if it occurred on the expected day and no 100% cornified read was seen. When cycles were aligned by the first cornified peak they were mostly regular during the first round of swabbing. A few rats drifted during the break, almost always by one day (Figure 5a). Realigning the cycles to the first cornified peak in the second tracking round showed that consecutive cycles are overwhelmingly regular (Figure 5b). Over the entire experiment, the majority of cycles lasted four days (Figure 5c). A total of 237 cycles were followed, of which 211 were four-day, 13 were five-day, and 5 were three-day.

**Figure 5.**
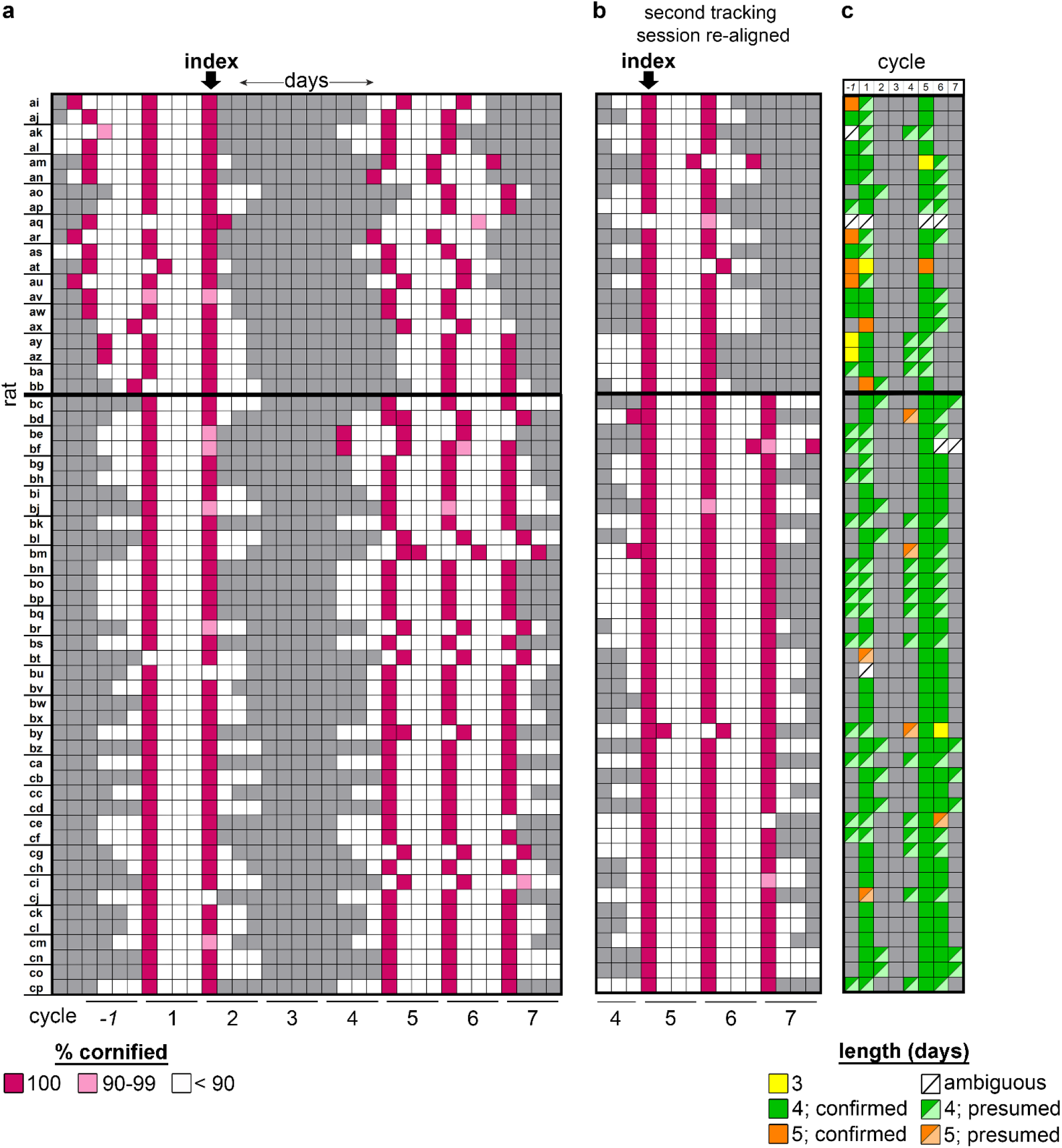
Track-by-Day in a large cohort across four weeks. (a) Daily smears were collected over two rounds separated by a break. Two groups (n=20 and n=40) were tracked with slightly different collection and break schedules. Smears were categorized as either fully cornified or not, although in a few instances 90-99% cornification presented in the smear when a peak was expected (light pink). Cycles were indexed to the final cornified peak before the break. Gray boxes indicate days when no smear was collected, and lines on the y-axis show cage-mate pairs. (b) The second tracking round realigned using its first cornified peak as the index day. (c) Across 60 animals, 240 full cycles were tracked: the majority were four-day (211 of 237, 89.0%), while the rest were either three-day (5 of 237, 2.1%), five-day (13 of 237, 5.5%), or ambiguous (8 of 237, 3.4%). Importantly, by the second tracking session (PND 65), the incidence of four-day cycles increased (125 of 136, 91.9%), therefore 97% of females (58 of 60) were regularly cycling. Gray boxes in (c) refer to alignment resulting in an incomplete cycle, i.e., less than four or five days of cytology data.

The behavior experiment began 1 – 4 days after the final day of smear collection (Supplemental Figure 5-1). The final cycle was used to assign females pseudorandomly to a paired fear learning or unpaired control protocol to ensure representation of cycle days in each experimental group. A few females were assigned to a third tone-only group whose behavior data were not analyzed. Males were match-handled and balanced across experimental groups as well. Training and long-term memory testing were performed four days apart so that they would occur on the same day of a (four-day) cycle (Figure 6a). We intentionally avoided training and testing on Day 1 for two reasons. First, our twice-daily swabbing experiments found that the cornified peak is most frequently observed in the late dark period and early light period (Figure 4a, c), which would mean that our dark period Day 1 smears are taken during the pre-ovulatory drop in progesterone (Figure 1a). This drop is rapid, and even with experiments performed within a 4-hour time window, there could be dramatic differences in hormone levels within a Day 1 experimental group. Second, five-day cycles are expected to have two consecutive days of high 17β-estradiol (Figure 1a) and potentially two days of peak cornified reads (Figure 4a). In this case, a Day 1 smear and a Day 5 smear are indistinguishable without the following day’s smear to confirm a four- or five-day cycle. A final smear was collected from each female after the long-term memory test, and although we attempted to predict and avoid Day 1, four final smears predicted to be Day 4 or Day 2 showed Day 1 cytology. These rats were excluded from behavior analysis.

**Figure 6.**
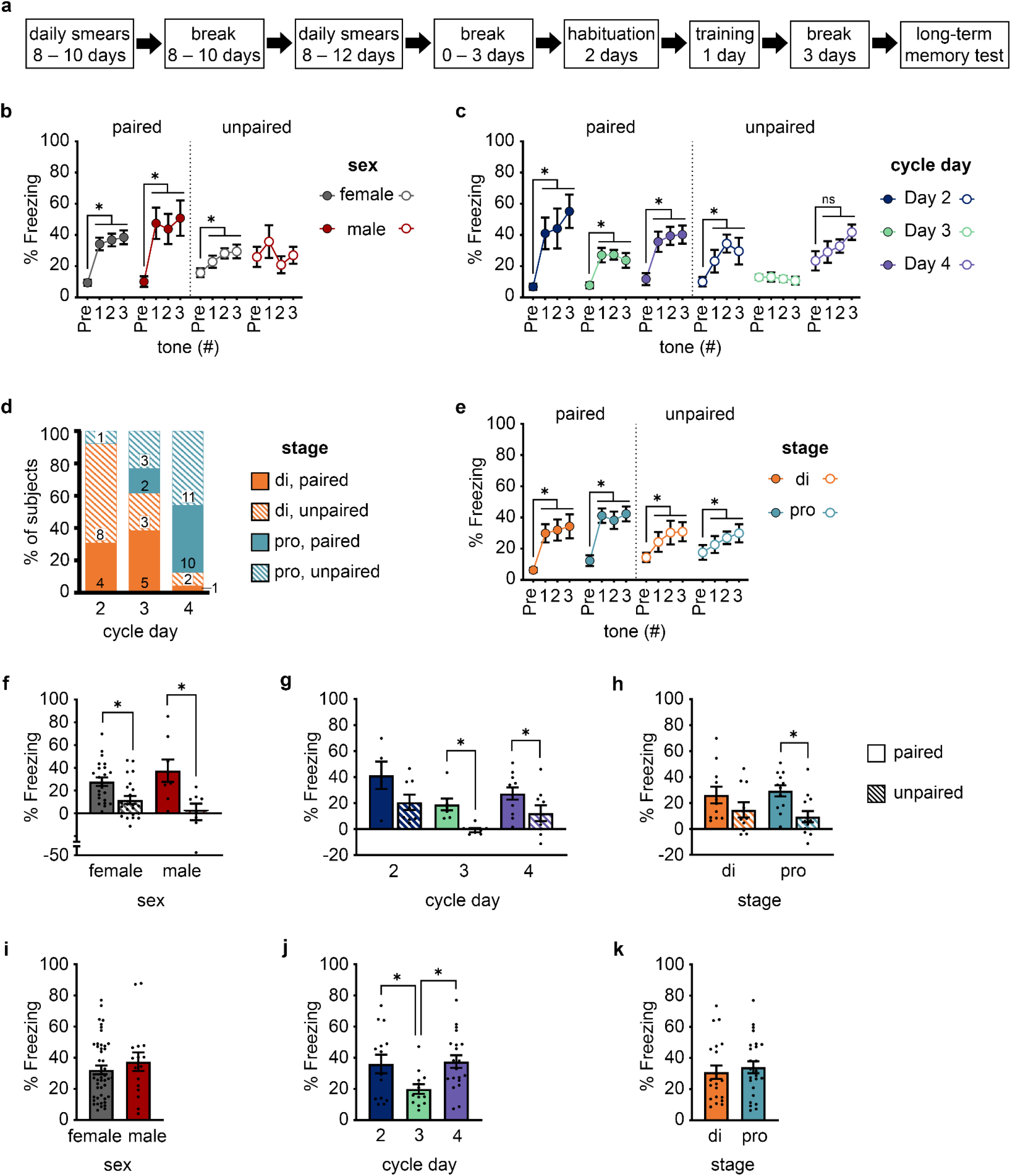
Track-by-Day detects estrous cycle effects on Pavlovian fear conditioning. (a) Experiment timeline. (b) Freezing before the first tone presentation (Pre) and during the three test tones during the long-term memory test in males and females. Freezing to the tone relative to the pre-tone period was higher in both sexes after paired training, and in females after paired training. (c) Tone freezing was higher than pre-tone after paired training on all cycle days, and after unpaired training on Day 2. (d) Distribution of female subjects by traditional cytology stage and cycle day. (e) Tone freezing was higher in diestrus and proestrus females regardless of training. (f-h) The difference between tone and pre-tone freezing was greater in the paired group versus the unpaired group in both sexes (f), on Day 3 and 4, but not Day 2 (g), and during proestrus but not diestrus (h). (i) Total freezing during the tone for both training groups was not different between males and females. (j-k) Among females, total freezing during the tone in both training groups was lower on Day 3 than on Days 2 and 4 (j), but did not differ between diestrus and proestrus (k). * p<0.05

During the long-term memory test, both males and females in the paired training group showed higher freezing to the tones relative to the pre-tone period. Females, but not males, in the unpaired group also showed higher freezing to the tone, indicating non-associative effects of training (Figure 6b). Females on all cycle days showed increased freezing to the tone after paired training, but only unpaired subjects on Day 2 showed non-associative freezing (Figure 6c). To compare Track-by-Day to traditional staging, a blinded, experienced experimenter used conventional cytology guidelines to classify the final smears for females that underwent behavior procedures as diestrus or proestrus. Stages markedly overlap, with Day 2 consisting mainly of diestrus reads and Days 3 and 4 of both diestrus and proestrus (Figure 6d). As in the previous dark-period experiment, leukocytes were very scarce and thus no animals were assigned to the metestrus stage. When the female data were analyzed by stage, no difference was seen between diestrus and proestrus subjects (Figure 6e). Relatively high freezing was seen in the unpaired group overall. Another cued fear conditioning study conducted during the dark period reported similar results^46^, suggesting that context generalization may be higher in the dark period. To isolate tone responses, pre-tone freezing was subtracted from tone freezing. Both males and females had significantly higher freezing to the tone in the paired group (Figure 6f), confirming associative learning to the tone. Among females, freezing in the paired and unpaired groups did not differ in subjects on Day 2 (Figure 6g) or those classified as diestrus (Figure 6h), consistent with the non-associative tone freezing in these unpaired groups. Overall freezing did not differ between males and females (Figure 6i) but was significantly lower on Day 3 than on Days 2 and 4 (Figure 6j). Diestrus and proestrus did not differ (Figure 6k), meaning that Track-by-Day was more sensitive in detecting a cycle-related change in behavior.

Training was performed using the same shock parameters for all animals, but shock sensitivity has been reported to be affected by sex and hormonal state ^47–49^. Because lower freezing on Day 3 and non-associative sensitization on Day 2 could be explained by lower and higher shock sensitivity, respectively, we evaluated shock sensitivity in a separate cohort of female and male rats. Females displayed higher sensitivity than males, but there were no differences between cycle days (Supplemental Figure 6-2).

### Additional cell types and overall cell numbers associated with Day 2

The final smears from the behavior experiment were stained with Shorr stain, which reveals cytology in finer detail than H&E and allowed five cell types to be quantified instead of three. Cornified cells largely appear bright orange with Shorr stain, and the full cornification of Day 1 is easily detected (Figure 7a). Nucleated epithelials show clear contrast between the nucleus and cytoplasm and high variability on Days 2-4 (Figure 7b). Leukocytes were very rare in these smears (Figure 7c), consistent with our earlier H&E-stained smears taken during the dark period (Supplemental Figure 4-3). Non-nucleated epithelial cells are difficult to differentiate with H&E staining but are very clear with Shorr stain. Acidophilic non-nucleated epithelials were very sparse (Figure 7d). In contrast, basophilic non-nucleated epithelials, recognized by bright cyan staining in crisp round cells, appeared frequently on Days 3 and 4 but rarely on Day 2 (Figure 7e). Independent of the specific cell types, the absolute number of cells per smear was higher on Day 2 than on Days 3 and 4, but there were no differences between stages (Supplemental Figure 7-1). Based on gradual changes in cytology, some smears were classified as late estrus (EL), late diestrus (DL), or late proestrus (PL). Similar to the canonical stages (Figure 6e), transitional stages progressed across days (Supplemental Figure 7-1). Number of cells per smear were not different when compared by stages of the estrous cycle (Supplemental Figure 7-2). Although not as unmistakable as the fully-cornified cytology of Day 1, these two features could be helpful in confirming Day 2 by large cell numbers or a paucity of basophilic non-nucleated epithelial cells.

**Figure 7.**
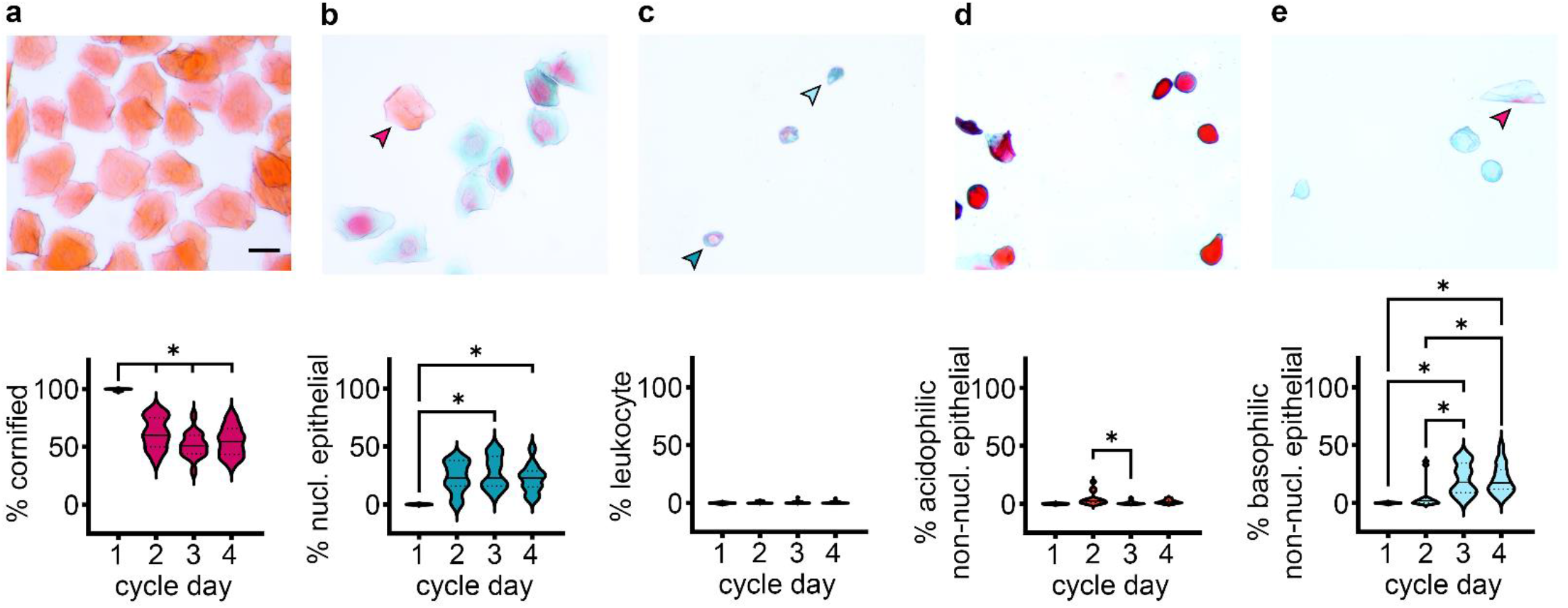
Quantitative cytology using Shorr stain. (a-e) Appearance of Shorr-stained cells and their percentage on the post-testing smears from the 60 rats in the behavior experiment: (a) cornified; (b) nucleated epithelial cells, with a stray cornified cell (arrow); (c) a leukocyte near a small nucleated epithelial cell (dark arrow) and a basophilic non-nucleated epithelial cell (light arrow); (d) acidophilic non-nucleated epithelial cells; (e) basophilic non-nucleated epithelial cells with a stray basophilic cornified cell (arrow), distinguished by its shape. Scale: 25 µm. * p<0.05

### Cycle days are more predictive of uterine histology than traditional stages

Unlike the vaginal epithelium, uterine tissue is sensitive to hormone changes throughout the estrous cycle^17,50^. If the hormone cycle is temporally stereotyped, uterine changes should correlate better with cycle days than with cytology stages. Immediately after collection of the final post-long-term memory test smears, all rats were perfused with fixative and the uterine tissue was harvested for histology. Gross examination of uterine sections across days (Figure 8a) revealed a dramatic dilation of the lumen in all five rats with Day 1/estrus cytology (Figure 8b). The only other rat with this uterine morphology was the one that did not have any four- or five-day cycles and did not have a cornified peak during the final round of smear collection (Figure 5c); this combination of persistent non-estrus cytology and histology confirms pseudopregnancy.

**Figure 8.**
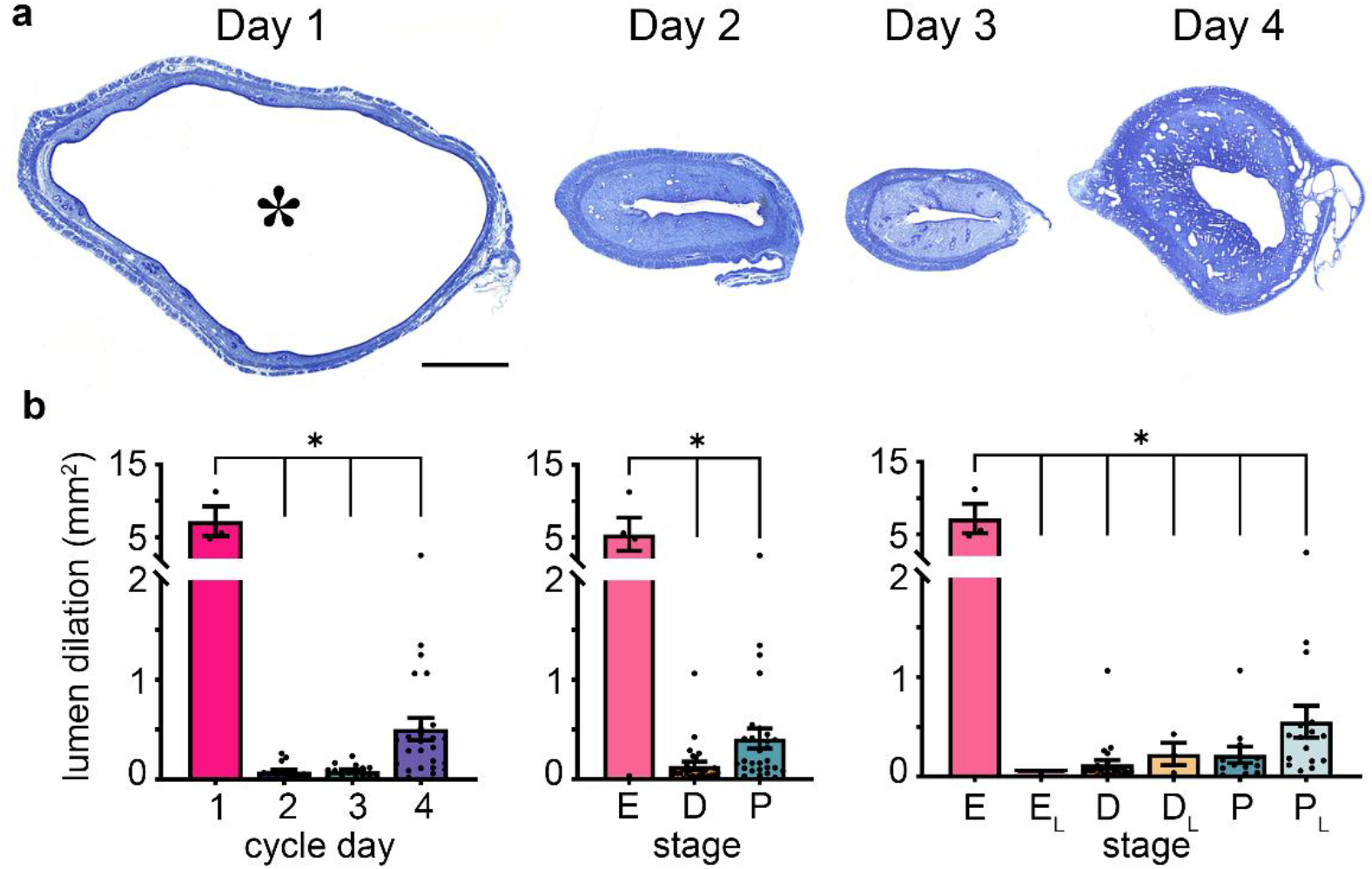
Gross uterine histology across the cycle. (a) Representative uterine cross sections from each cycle day. The lumen of the Day 1 sample is indicated with a large asterisk. (b) Extreme lumen dilation differentiates Day 1/estrus histology from the rest of the cycle. Scale: 1 µm. * p<0.00005.

Finer histological analysis revealed features associated with each of Days 2 – 4. Mitotic figures in the luminal epithelium (Figure 9a) were usually present on Days 3 and 4 but never seen on Day 2. In contrast,luminal mitosis did not differ between diestrus and proestrus or among the transitional stages (Figure 9b). The presence of glands in the endometrium (Figure 9c) was also lower on Day 2 versus Days 3 and 4 but was constant between stages (Figure 9d). Mitotic figures were present in the endometrial glands (Figure 9e), and their density distinguished Day 3 from the other days but did not differ between stages (Figure 9f). Finally, the cross-sectional area of the endometrium (Figure 9g) was greater on Day 4 than Days 2 and 3 (Figure 9h). This was the only histological measure that differed between stages, being greater in proestrus than diestrus, and greater in late proestrus than early proestrus. Given that two-thirds of proestrus and all late proestrus cases occurred on Day 4 (Supplemental Figure 7-1), this is not surprising.

**Figure 9.**
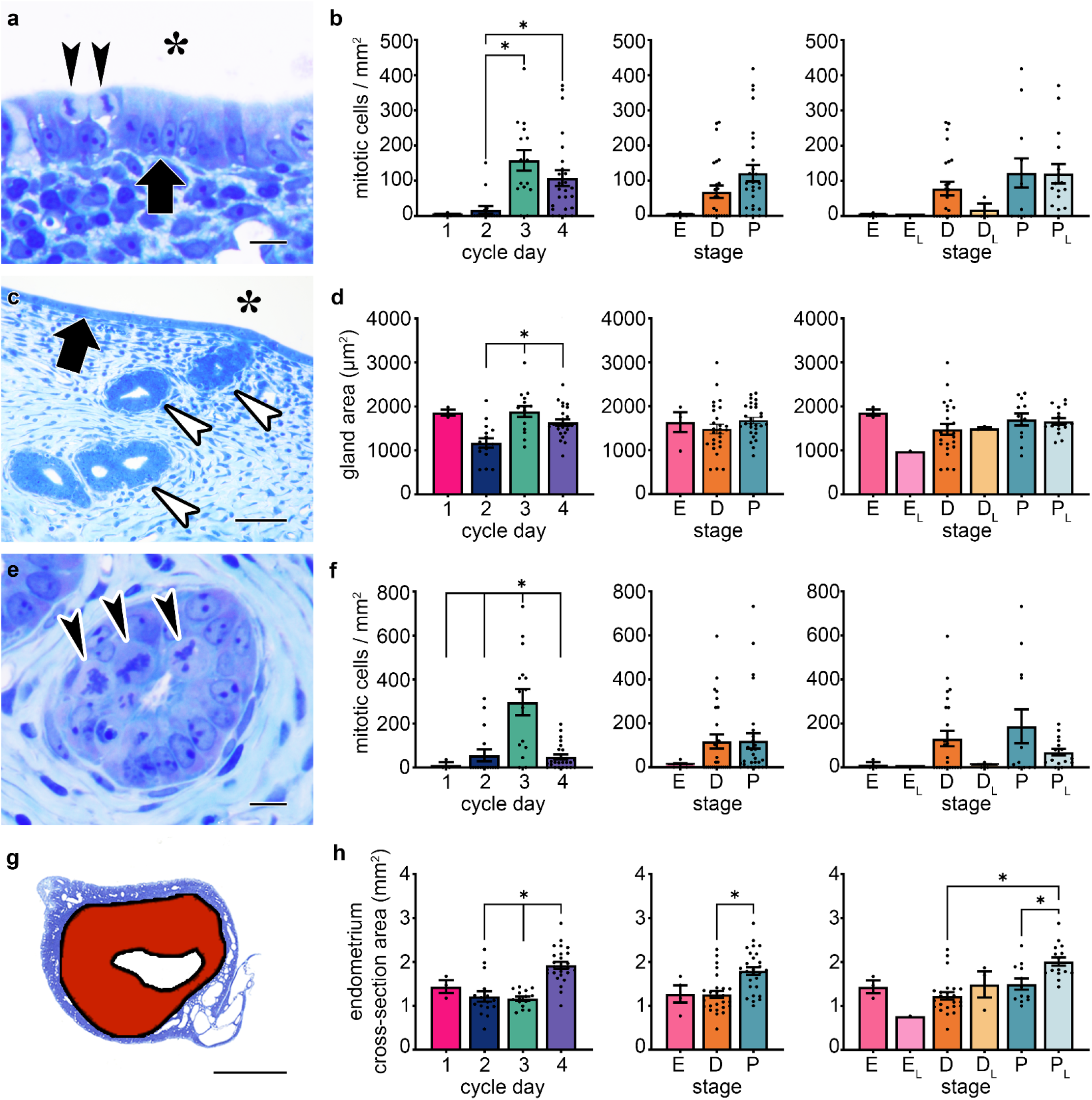
Cycle days correlate with changes in uterine histology. (a) Mitotic cells (arrowheads) in the luminal epithelium (arrow), which lines the lumen (large asterisk). (b) There were fewer luminal mitotic cells on Day 2 versus Days 3 and 4, but no differences between the stages. (c) Endometrial glands (arrowheads) below the lumen (asterisk) and luminal epithelium (arrow). (d) Endometrial glands covered less area on Day 2 versus Days 3 and 4, but the stages did not differ. (e) Mitotic cells (arrowheads) in an endometrial gland. (f) There were more mitotic cells in endometrial glands on Day 3 versus other days, but no differences between stages. (g) Uterine cross section with endometrium shown by red fill. (h) The cross-sectional area of the endometrium is higher on Day 4 than on Days 2 and 3, and is also higher in proestrus than in diestrus and in late proestrus versus early proestrus and diestrus. Scale bar = 25 µm (a, c, e), 1 µm (g). E: estrus, E_L_: late estrus, D: diestrus, D_L_: late diestrus, P: proestrus, P_L_: late proestrus. * p<0.05

## Discussion

The lack of straightforward, accessible methods for tracking the rodent estrous cycle is a barrier to progress in understanding sex differences in the brain. There is disagreement in the literature as to whether and how the estrous cycle affects fear and anxiety-related behaviors and brain functions^12,13^, which validates the perception of the cycle as a source of uncontrollable variability. This concern is reflected not only in the ongoing exclusion of female subjects from behavioral research^3,4,51^, but also in the common practice of ovariectomizing female subjects^14^. The problem is less that the cycle may introduce variability, and more that the cycle is so difficult to monitor that it is preferable to alter subjects with an invasive procedure and study them under the influence of synthetic hormones. Traditional tracking methods are subjective and require expert training, which deters widespread use, and worse, they are not standardized. One popular guide to staging, written by toxicologists, recommends that “individual laboratories define their respective processes” for staging^16^. This approach would be effective for evaluating whether a given toxin disrupts cycling, for example, but is likely to lead to irreproducible results if used to compare behaviors between points in the cycle. In this study, we describe a simple, straightforward method for tracking the estrous cycle that provides a universal reporting framework. Using Track-by-Day, we show that the estrous cycle is highly predictable and regular across subjects and conditions. We demonstrate that the method outperforms traditional staging in detecting behavioral effects, and correlation with uterine histology confirms its greater physiological relevance. Our data counter perceptions that tracking the cycle requires expert knowledge and that irregular cycles are common^15,16,27^.

The first description of changes in vaginal cytology during the rat estrous cycle was published 100 years ago^24^. Cytology was divided into stages according to a four-stage model of mammalian reproduction, which itself was based on observations of reproductive physiology and behavior across many species ^22^. It was not until decades later that the hormone cycle was measured in detail and the effects of steroid hormones on the vaginal epithelium were investigated. Several studies have monitored serum levels of ovulation-related hormones across the rat estrous cycle^19–21^, and two features of the cycle have clear practical implications for cycle tracking. First, hormone levels follow a stereotyped pattern that is entrained to the light-dark cycle, meaning that cycle progression can be inferred if a single time point can be identified. Second, the major hormonal changes – a surge in 17β-estradiol followed by sharp, nearly simultaneous peaks of progesterone, luteinizing hormone, follicle-stimulating hormone, and prolactin – occur in the 24 hours preceding ovulation, and the traditional cytology stages do not correspond to hormone states. Since stage assignments are effectively arbitrary with respect to hormones (with the exception of estrus), inconsistency between studies is unsurprising. Dividing subjects on Day 2 between metestrus and diestrus creates two groups with essentially identical hormone levels, whereas the cytology does not reflect the rapid hormonal changes on Day 4 in real time and subjects assigned to diestrus or proestrus on Day 4 are unlikely to differ systematically.

Our data show that the most reliably cyclic feature of cytology is the cornified peak and that variability in cytology increases as the cycle progresses, consistent with the relationship between cytology and the hormone peaks becoming more indirect. Our observations are also consistent with cytology guidelines that describe nuanced stage transitions and broad ranges for stage durations. Timing by the cornified peak alone, we found that the vast majority of cycles lasted four days and that almost all rats had regular cycles. This counters the common perception that cycles are often irregular, or, as stated in the title of a recent guide to staging^15^, that there is “no such thing as a normal cycle.” If a normal cycle requires each of the traditional stages to appear in order or for consistent durations, most of our rats would be considered irregular. Since it is common to exclude subjects without regular cycles and there is no agreed-upon definition of regular, differences in exclusion criteria could explain some conflicting results in the literature.

Uterine histology is a more direct reflection of cycle progression than vaginal cytology, as uterine tissue responds more rapidly to 17β-estradiol, and there is evidence that unlike in the vaginal epithelium, the response is dose-dependent^36,52^. Descriptions of changes in uterine histology across the cycle are generally consistent: the lumen dilates on the day of ovulation, accompanied by necrosis of the epithelium and glands^17,53,54^. Water is then expelled from the lumen, and necrosis slows while mitosis resumes, restoring the mature epithelial and gland structures. Our data reflect this temporal progression. The cornified peak in vaginal cytology corresponded 1:1 with profound lumen dilation, and mitosis followed by uterine enlargement differentiated the ensuing days. Only uterine size differed between traditional stages, and variability was higher between stages than between days.

Similarly, comparing across cycle days detected cycle-related differences in cued Pavlovian fear conditioning that were not apparent between cytology stages. Studies examining the effects of sex and estrous stage on Pavlovian conditioning are inconsistent, with some reporting no differences and others reporting differences in opposing directions^12–14^. Consistent with other studies performed in the dark period^46,55,56^, we found that subjects of both sexes and all estrous stages and days displayed tone-evoked freezing after paired training. Like another study^55^, we looked for darting^57^ but found that it was unusual and not systematic, suggesting that this behavior may be more common during the light period. Unpaired training produced non-associative freezing in females on Day 2, while Day 3 subjects displayed low freezing regardless of training.

When the cycle was not accounted for at all and when traditional staging was used, non-associative freezing appeared to occur in females generally as opposed to in a subset of subjects. Meanwhile, the freezing effect on Day 3 was completely obscured when subjects were grouped by stage. Neither non-associative freezing nor lowered freezing can be attributed to differences in shock sensitivity, which did not change across the cycle. Non-associative freezing is likely due to pseudoconditioning, which is mechanistically distinct from associative learning and thought to be a non-specific arousal effect^58–60^. Interestingly, a recent study of male and female rats in a naturalistic foraging setting found that a tone stimulus evoked non-associative freezing but not Pavlovian conditioning^61^. Pseudoconditioning in Day 2 females could thus represent ethologically-relevant vigilance or defensive behavior that is reduced later in the cycle. Lowered freezing on Day 3 does not indicate impaired learning, as associative freezing to the tone was preserved, but instead a difference in behavioral expression.

Grouping by cycle day in the dark period yields groups with presumed low hormones (Day 2), high progesterone (Day 3), or high 17β-estradiol (Day 4), so unlike staging, Track-by-Day allows interpretation in the context of hormone levels. Endogenous and exogenous progesterone and 17β-estradiol have been reported to be anxiolytic in females^14^. In the elevated plus maze, progesterone is associated with more open arm entries as well as more open arm time in the elevated plus maze^62–64^, whereas acutely high 17β-estradiol has been associated with more open arm time but not more entries^65,66^. Although freezing in a Pavlovian conditioning paradigm is not considered a measure of anxiety, these effects are interesting in the context of our data. If both steroids reduce anxiety responses, the non-associative freezing on Day 2 could reflect higher vigilance in the low-hormone state. If progesterone drives more active behavior in response to threats than 17β-estradiol, that could explain the overall low freezing on Day 3.

Overall, tracking data from 328 cycles in 94 rats demonstrated that cycle length is regular across subjects, cycles, and conditions. Most cycles (85%) lasted four days, and only four rats failed to have any four-day cycles. One of these had no discernable cycles and was confirmed to be pseudopregnant by uterine histology, while the other three had five-day cycles. Most five-day cycles were seen in rats that also had four-day cycles, however, meaning that five-day cycles occur sporadically as opposed to being purely characteristic of individuals. Synchrony between cage-mate pairs occurred at chance levels, consistent with a quantitative study of cycle timing in group-housed rats^40^. The stress of daily blood draws did not disrupt cycle length, consistent with a report of four-day cycle length persisting during daily restraint stress^55^. It is important to note that we collected all of our data on young adult Sprague-Dawley rats, and the cycle may not manifest identically at other ages or in other strains. Cycle length and regularity are reported to vary in rats younger than one month and older than ten months^67–69^, so our results should not be assumed to apply across the entire life cycle.

In contrast to traditional staging, the Track-by-Day method does not require specialized training in cytology interpretation, making it accessible to a wide community of researchers. It also inherently facilitates standardization across laboratories; regardless of whether traditional stages are given, studies that report the cycle day and time of data collection can be easily compared. It is important to note that in classic endocrinology literature^19–21^ the names of the four stages clearly refer to days as opposed to cytology, and at least one neuroscience group follows this practice^55,56^. Because of the longstanding and extensively reinforced association between the stage names and cytology states of variable duration, we propose a new practice of referring explicitly to cycle days as opposed to changing the use of staging terminology. By making a break with the stage names and their association with complex tracking methods, we hope that the Track-by-Day method will encourage more researchers to work with female subjects and improve our understanding of how the estrous cycle does, and doesn’t, influence behavior.

**Recommendations for the Track-by-Day Method**

<u>Planning:</u> Track smears for twelve consecutive days before beginning an experiment. This will acclimate the subjects (and experimenters) to the procedure and provide two full cycles of data.

<u>Sample collection:</u> Collect smears once per day, ideally between the mid-dark period and mid-light period. Contrast smears with H&E or Shorr stain.

<u>Interpretation:</u> Identify peak smears containing only cornified cells, especially in clumps and sheets.

<u>Tracking:</u> Define fully-cornified smears as Day 1, and assign non-peak smears to the corresponding cycle days. If two consecutive peak cornified smears are observed, the final smear before a non-peak smear is Day 1. This ensures that the cycle is timed from the day of ovulation.

<u>Reporting:</u> Subjects can be grouped for analysis by cycle days instead of or in addition to traditional stages, but whether or not stages are used, the cycle day relative to the cornified peak and the exact time that experiments were performed should be reported to allow standardized comparisons between studies.

## Materials and Methods

### Subjects

Subjects were adult female and male Sprague-Dawley rats (Hilltop Lab Animals Inc., Scottsdale, PA) pair-housed with *ad libitum* food and water. After arrival from the vendor, animals were acclimated to the housing room for a minimum of five days before experiments began, or a minimum of ten days for those housed with a reverse light cycle. All animal procedures were approved by the Institutional Animal Care and Use Committee at the University of Connecticut.

### Collection of vaginal smears

Animals were gently restrained and a saline-dipped cotton-tipped applicator (2 mm diameter, cotton tip length 1 cm) was inserted into the vagina to the depth of the cotton tip. Swabs were gently rolled out onto a gelatin-subbed microscope slide and the smears were allowed to dry before any staining. Male rats were subjected to identical handling, and individual experimenters did not handle males and females on the same day.

### Staining, imaging, and analysis of vaginal smears

For cresyl violet staining, slides were dipped in 0.1% aqueous cresyl violet (Millipore Sigma, Inc.) for 5 minutes, then rinsed in water. H&E (Vector Labs, Inc.) and Shorr (Millipore Sigma, Inc.) stains were used according to the manufacturers’ instructions. Smears used for quantification were stained with H&E except for the terminal smears taken after the behavior experiment, which were Shorr stained. Stained slides were dehydrated in ethanol, cleared in toluene, and coverslipped with DPX mounting medium. Smears were imaged at 10X using either a Keyence BZ-X700 microscope or on a compound microscope with a Canon EOS 800D digital camera. Reconstruct software ^70^ was used for cell quantification. A counting frame of 285.75 mm × 285.75 mm was applied to a representative area of each image, and all cells within the counting frame were counted and classified. Three separate images were analyzed per smear by at least two blinded experimenters.

### Impedance measurement

To construct the probe, two gold pins (3 cm long by 1 mm wide) were soldered to copper wire with lead-free silver solder, and heat shrink was used to secure them in a parallel position 3 mm apart. For impedance readings, the probe was attached to a standard multi-meter and inserted to a depth of ∼2 mm so that the pins were in contact with the dorsal and ventral vaginal wall. Impedance readings were taken prior to swabbing for cytology smears, and the probe was sanitized with 100% ethanol and allowed to dry before and after each reading.

### Pavlovian fear conditioning

All behavioral experiments were conducted in the dark period, 3–7 hours after lights-off. The behavioral apparatus, training, and testing protocols were identical to those used in Ostroff et al. (2010)^71^ with the exception that the conditioning chambers were unlit during all procedures. Briefly, animals were habituated to the conditioning chamber (Coulbourn Instruments) for 30 minutes on each of two consecutive days before training. Tones (30s, 5kHz, 80dB) were delivered through a speaker in the chamber and shocks (1s, 0.7mA) were delivered through a grid floor. Paired training consisted of five tones co-terminating in footshocks over a single 32.5 minute training session, and unpaired training consisted of five non-overlapping tones and shocks. Testing consisted of three tones presented in the same chamber, with the context modified by the addition of a smooth opaque acrylic floor and the scent of peppermint. Cage-mate pairs were trained and tested at the same time using the same protocol, and because pairs of females did not synchronize their cycles it was impossible to collect equal numbers of each stage. Freezing during the tones was scored by three experimenters blinded to condition, sex, and stage, and the scores were averaged for analysis.

### Shock sensitivity

Adult (PND 90) female (n=26) and male (n=8) rats were habituated to the reverse light-cycle housing room for at least two weeks. Females were tracked for four cycles leading up to the experiment and testing was divided across four days to ensure that all cycle days were represented. Males were match-handled and run separately after the females. Shock sensitivity was performed with the same equipment used for Pavlovian conditioning during the mid-dark cycle. Rats were placed singly in a testing box and a series of ten shocks were presented at 40s intervals. Shock intensity began at 0.1 mA and increased in steps of 0.1 mA, so that the final shock was 1 mA. Cage-mate pairs were tested in parallel. The intensity of the first shock evoking a shuffle was recorded for each animal. Shuffles were defined as a fast, frenzied movement; some animals may shuffle in place by quickly moving all four paws.

### Collection of uterine tissue

One hour after long-term memory testing, rats were deeply anesthetized with chloral hydrate (750mg/kg i.p.) and perfused transcardially with 4% paraformaldehyde and 0.1% glutaraldehyde in 0.1M phosphate buffer (pH 7.4) at room temperature. The uterine horns were removed and post-fixed overnight in 4% paraformaldehyde, then stored in phosphate buffered saline with 0.01% sodium azide until further processing. Segments 0.5 to 1 cm long and roughly 1.5 to 2 cm away from the oviduct were dissected for embedding. Samples were dehydrated in ascending concentrations of ethanol, then infiltrated with a 1:5 mixture of methyl methacrylate and butyl methacrylate with 0.5% benzoin methyl ether. Samples were cured in gelatin capsules under UV light for 48 hours in a freeze-substitution unit (Leica). All dehydration, embedding, and curing were performed at 0ºC. The cured blocks were trimmed to expose a full cross section of the uterus, and 1µm sections were cut on an ultramicrotome (Leica). Sections were collected on gelatin-subbed slides, stained with 0.5% toluidine blue in 1% sodium borate for five minutes, and coverslipped with DPX. One representative cross-section was imaged per rat at 10X on a Keyence BZ-X700 microscope, and additional 40X images were taken on a compound microscope with a Canon EOS 800D digital camera.

### Analysis of uterine histology

Uterine area, endometrium area, myometrium area, luminal epithelium area, lumen dilation, and number and size of endometrial glands were measured on 10X images using Reconstruct software. Mitotic figures and tissue necrosis were quantified in 40X images. Mitotic figures were counted in the luminal epithelium and the epithelium of the endometrial glands and normalized to area. Each analysis was performed by at least two blinded experimenters.

### Statistics

Means of more than two groups were compared using one-way ANOVAs, and effects significant at p < 0.05 were followed with a Bonferroni post-hoc test. Unpaired t-tests were used when comparing two groups. Results of statistical comparisons are given in Supplemental Table 1.

## Acknowledgements

We are grateful to Maritza Abril, Zachary Deane, Jairo Orea, and Jerry Vasquez Jr. for help with some of the experiments, Teng Long for help with behavioral analysis, and Chris Cain, Michael Goard, Alex Jackson, John Redden, and Weiling Yin for helpful comments on the manuscript. This work was performed in part at the Bioscience Electron Microscopy Laboratory of the University of Connecticut.

---

**Figure 4 – Supplemental Figure 1.**
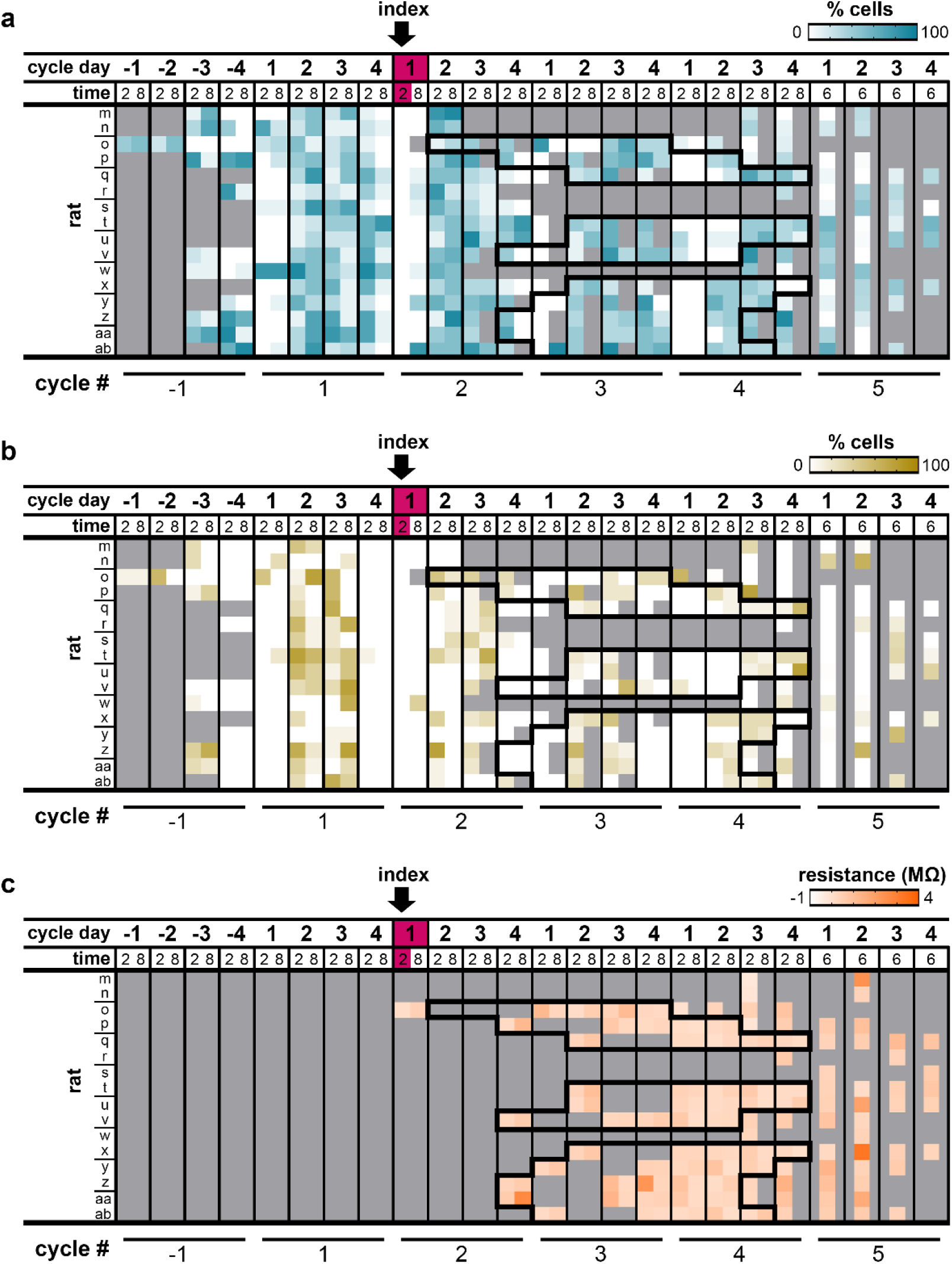
Quantified reads of non-cornified cell types in smears and impedance corresponding to Figure 4a. Black outline over measurements in cycles 2-4 indicates daily tail vein blood draws for specified rats. (a) Percent nucleated epithelial cells. (b) Percent leukocytes. (c) Vaginal impedance (MΩ) values. Lines for rats on y-axis indicate cage-mate pairs. Gray boxes indicate when smears/impedance were not collected.

**Figure 4 – Supplemental Figure 2.**
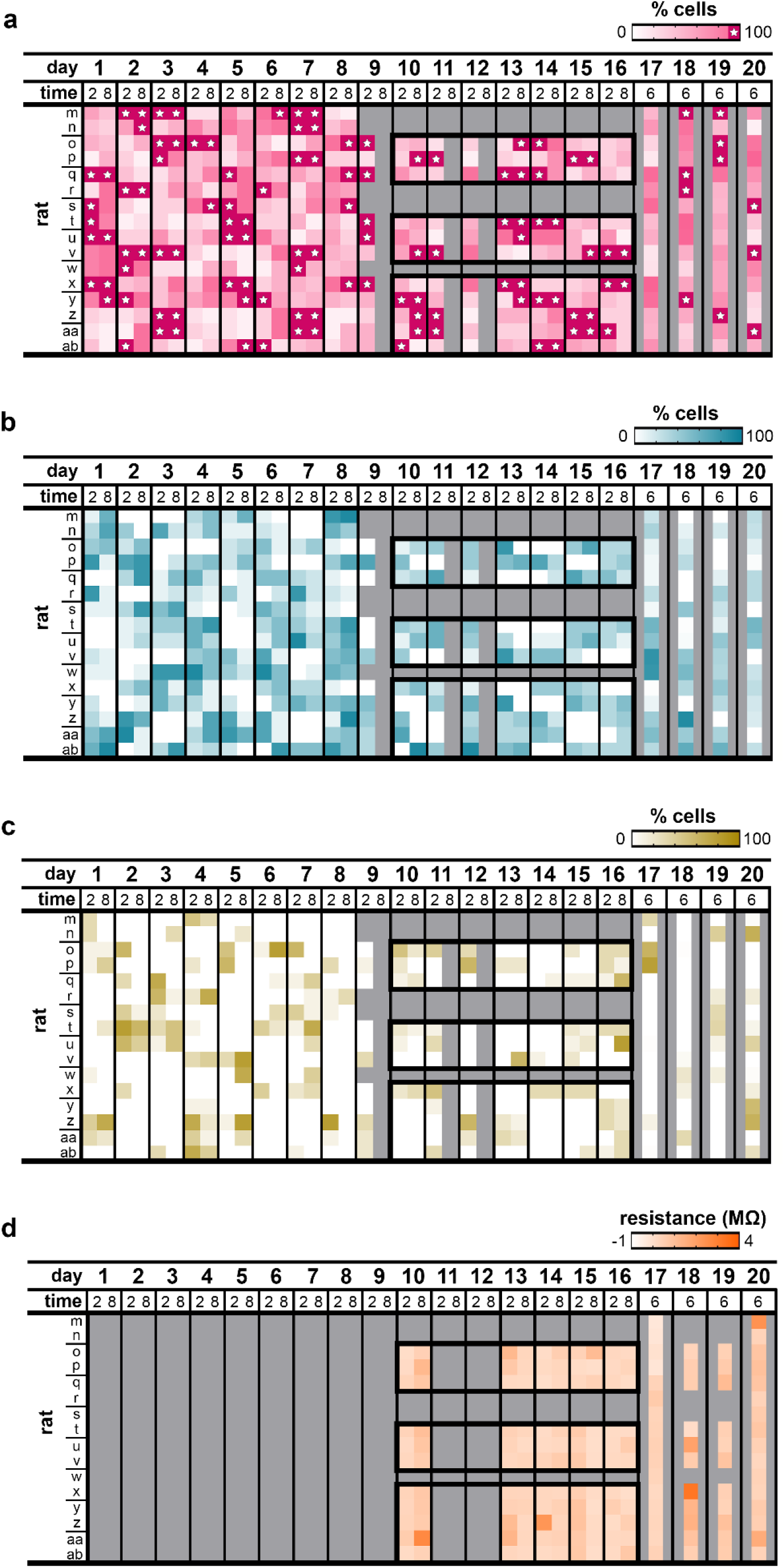
Unaligned data from Figure 4a and Supplemental Figure 4-1. (a) Percent cornified cells (data from Figure 4a). (b) Percent nucleated epithelial cells. (c) Percent leukocytes. (d) Vaginal impedance (MΩ) values. Lines for rats on y-axis indicate cage-mate pairs. Gray boxes indicate when smears/impedance were not collected.

**Figure 4 – Supplemental Figure 3.**
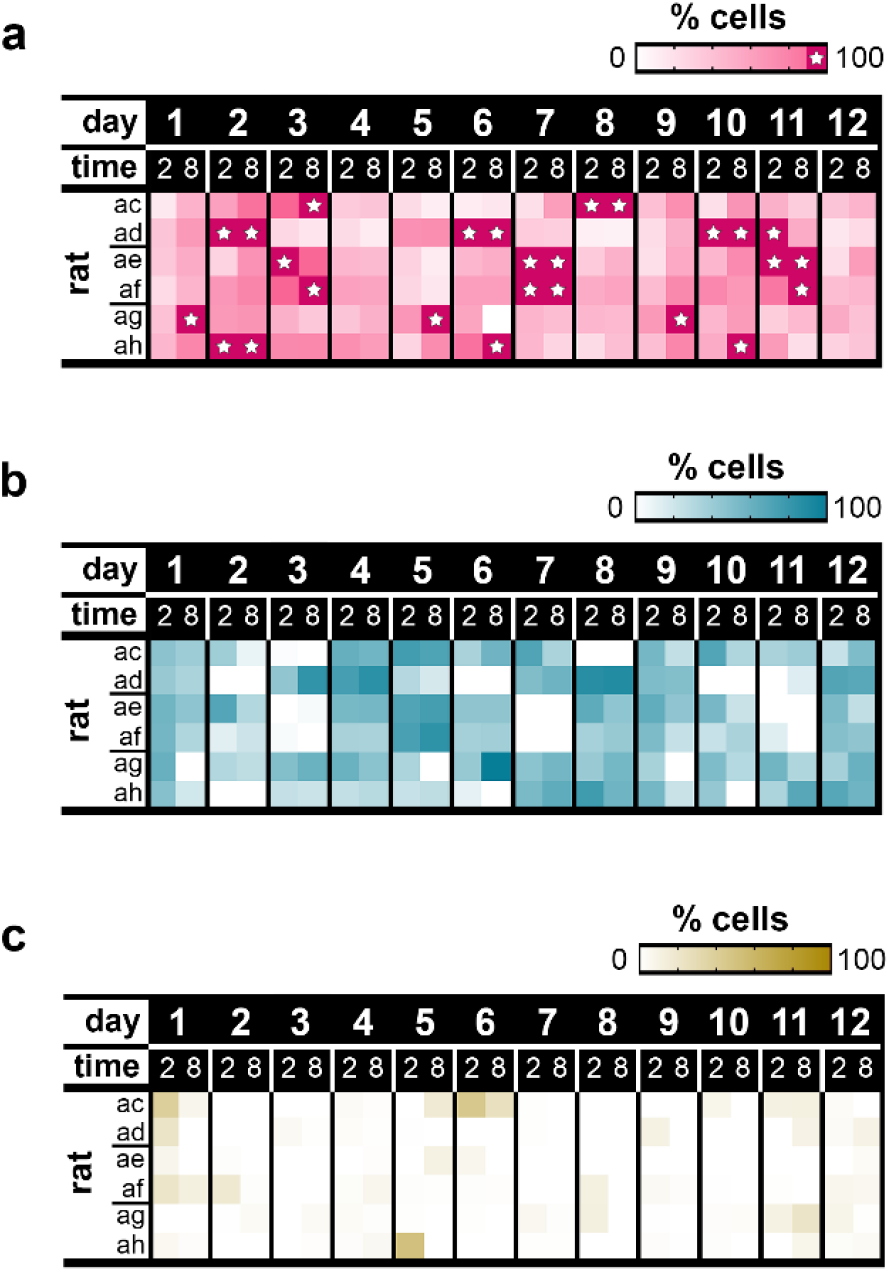
Unaligned data from Figure 4c and quantification of other cell types in the same smears. (a) Percent cornified cells (data from Figure 4c). (b) Percent nucleated epithelial cells. (c) Percent leukocytes. Lines for rats on y-axis indicate cage-mate pairs.

**Figure 4 – Supplemental Figure 4.**
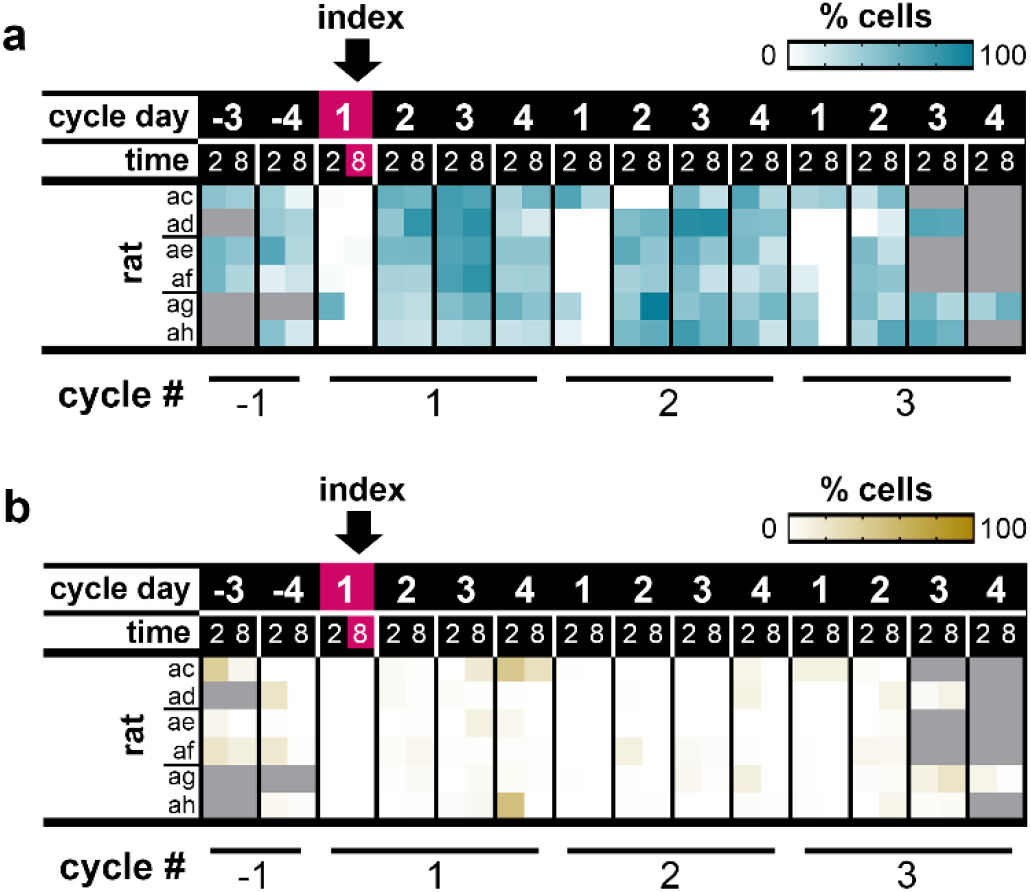
Aligned data from non-cornified cell types in smears corresponding to Figure 4c. (a) Percent nucleated epithelial cells (data from Supplemental Figure 4-3b). (b) Percent leukocytes (data from Supplemental Figure 4-3c). Lines for rats on y-axis indicate cage-mate pairs. Gray boxes indicate when data was not collected.

**Figure 6 – Supplemental Figure 1.**
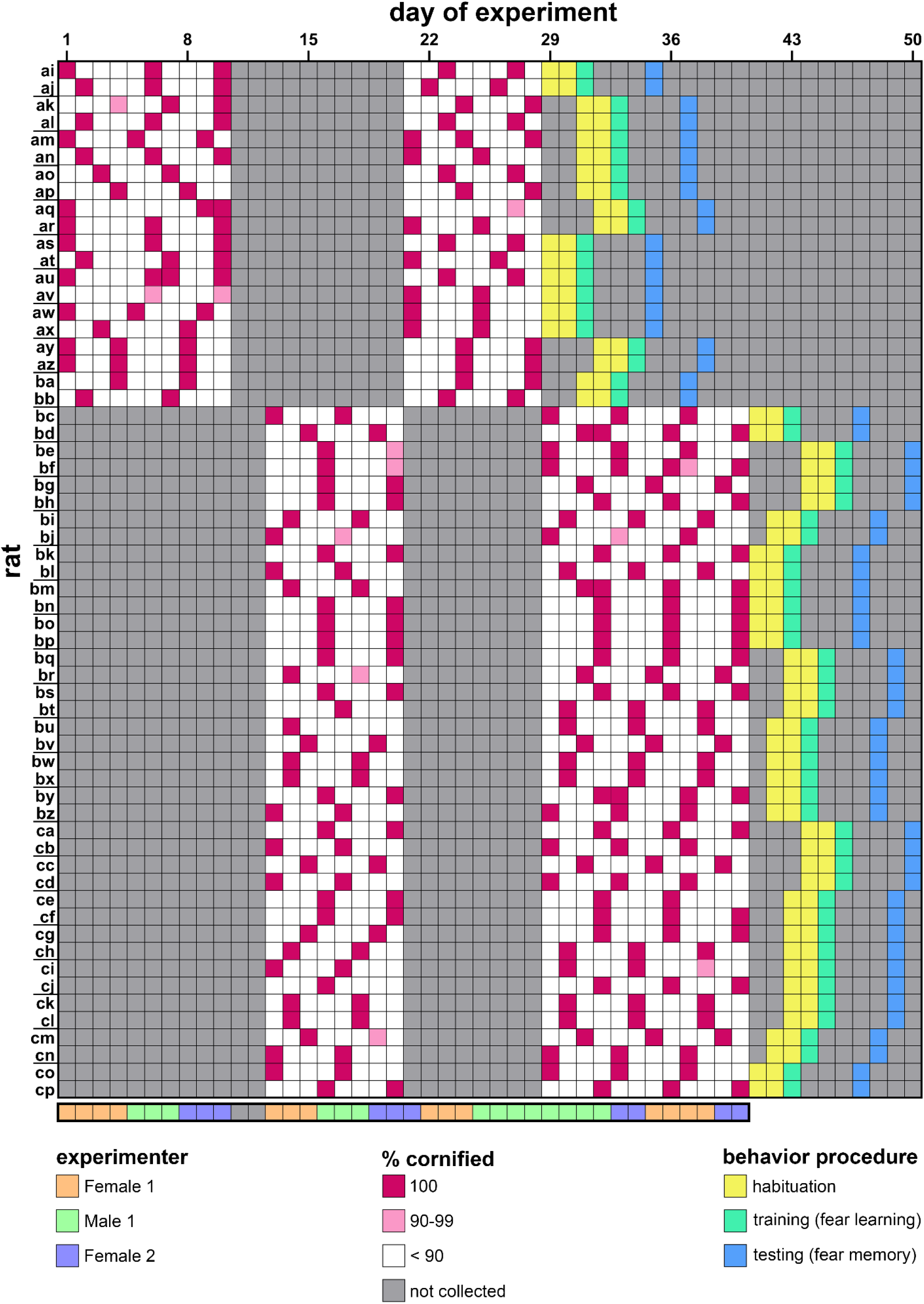
Unaligned tracking data from Figure 5a showing the full experiment over 50 calendar days, including the days of the behavior schedule in Figure 6a. Squares beneath tracking data indicate which of three experimenters (two female, one male) collected swabs on each day. Lines for rats on y-axis indicate cage-mate pairs. Gray boxes indicate when data was not collected.

**Figure 6 – Supplemental Figure 2.**
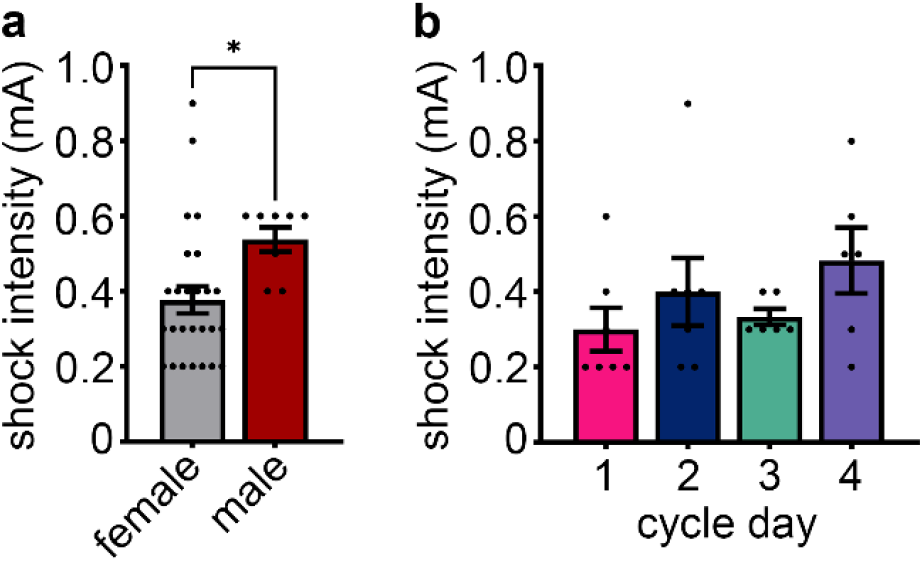
Sex differences in shock sensitivity, but no cycle day differences. (a) Females have a lower threshold to shuffle in response to a footshock than males * p < 0.05 (b) There was no effect of cycle day on shuffle threshold among females.

**Figure 7 – Supplemental Figure 1.**
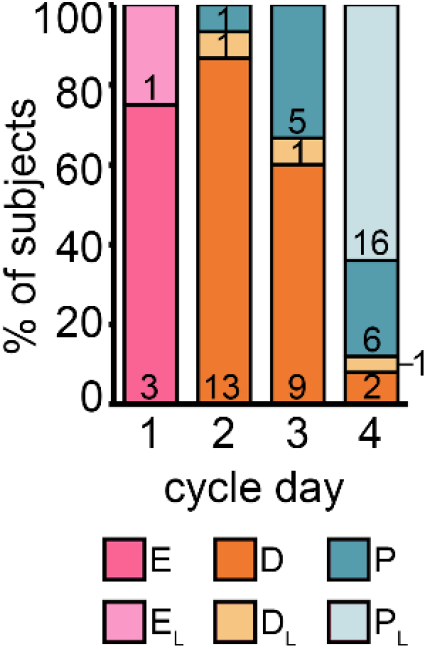
Shorr stain allows for subclassification of transitional estrous stages. Reassignment of females in Pavlovian conditioning experiment from cycle day to transitional cytology stage, and distribution of stages across cycle days. Numbers inside bars refer to quantity of rats per cytology stage. | E: estrus, E_L_: late estrus, D: diestrus, D_L_: late diestrus, P: proestrus, P_L_: late proestrus.

**Figure 7 – Supplemental Figure 2.**
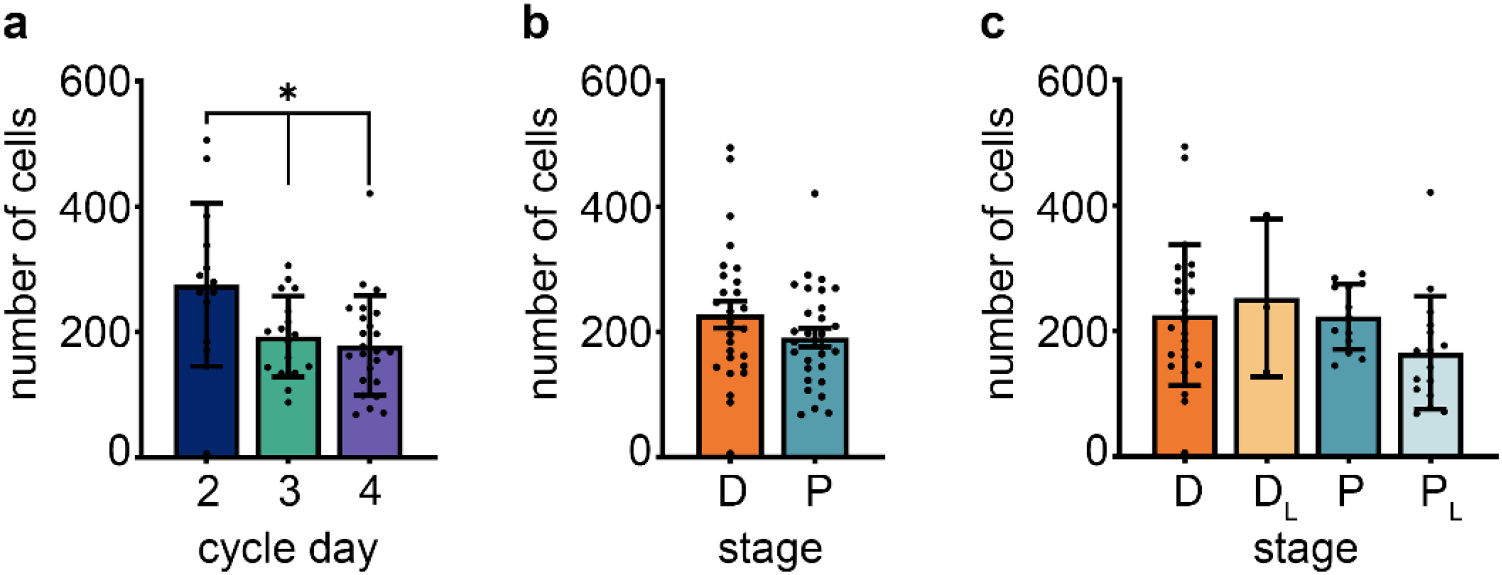
Number of vaginal cells present for each quantified smear. (a) There were more cells per smear on Day 2 than on Days 3 and 4. Day 1 was not quantified due to its usual composition of unquantifiable sheets and clumps. (b-c) No differences found when females are compared by major stages (b) or transitional stages (c). * p<0.05

## References

1. Pearse, R. V. & Young-Pearse, T. L. Lost in translational biology: Understanding sex differences to inform studies of diseases of the nervous system. Brain Research vol. 1722 146352 (2019).

2. Choleris, E., Galea, L. A. M., Sohrabji, F. & Frick, K. M. Sex differences in the brain: Implications for behavioral and biomedical research. Neuroscience and Biobehavioral Reviews vol. 85 126–145 (2018).

3. Mamlouk, G. M., Dorris, D. M., Barrett, L. R. & Meitzen, J. Sex bias and omission in neuroscience research is influenced by research model and journal, but not reported NIH funding. Frontiers in Neuroendocrinology vol. 57 (2020).

4. Hughes, R. N. Sex still matters: has the prevalence of male-only studies of drug effects on rodent behaviour changed during the past decade? Behav. Pharmacol. 30, 95–99 (2019).

5. Rubinow, D. R. & Schmidt, P. J. Sex differences and the neurobiology of affective disorders. Neuropsychopharmacology vol. 44 111–128 (2019).

6. Bangasser, D. A. & Cuarenta, A. Sex differences in anxiety and depression: circuits and mechanisms. Nat. Rev. Neurosci. 22, 674–684 (2021).

7. Shansky, R. M. Are hormones a “female problem” for animal research? Outdated gender stereotypes are influencing experimental design in laboratory animals. Science vol. 364 825–826 (2019).

8. Prendergast, B. J., Onishi, K. G. & Zucker, I. Female mice liberated for inclusion in neuroscience and biomedical research. Neurosci. Biobehav. Rev. 40, 1–5 (2014).

9. Becker, J. B., Prendergast, B. J. & Liang, J. W. Female rats are not more variable than male rats: A meta-analysis of neuroscience studies. Biol. Sex Differ. 7, (2016).

10. Zajitschek, S. R. K. et al. Sexual dimorphism in trait variability and its eco-evolutionary and statistical implications. Elife 9, 1–17 (2020).

11. Beery, A. K. Inclusion of females does not increase variability in rodent research studies. Current Opinion in Behavioral Sciences vol. 23 143–149 (2018).

12. Day, H. L. L. & Stevenson, C. W. The neurobiological basis of sex differences in learned fear and its inhibition. European Journal of Neuroscience vol. 52 2466–2486 (2020).

13. Lovick, T. A. & Zangrossi, H. Effect of Estrous Cycle on Behavior of Females in Rodent Tests of Anxiety. Frontiers in Psychiatry vol. 12 (2021).

14. Price, M. E. & McCool, B. A. Structural, functional, and behavioral significance of sex and gonadal hormones in the basolateral amygdala: A review of preclinical literature. Alcohol vol. 98 25–41 (2022).

15. Robert, H. et al. Rodent Estrous Cycle Monitoring utilizing Vaginal Lavage: No Such Thing As a Normal Cycle. J. Vis. Exp. 2021, (2021).

16. Cora, M. C., Kooistra, L. & Travlos, G. Vaginal Cytology of the Laboratory Rat and Mouse:Review and Criteria for the Staging of the Estrous Cycle Using Stained Vaginal Smears. Toxicol. Pathol. 43, 776–793 (2015).

17. Westwood, F. R. The Female Rat Reproductive Cycle: A Practical Histological Guide to Staging. Toxicologic Pathology vol. 36 375–384 (2008).

18. Frick, K. M. & Kim, J. Mechanisms underlying the rapid effects of estradiol and progesterone on hippocampal memory consolidation in female rodents. Horm. Behav. 104, 100–110 (2018).

19. Nequin, L. G., Alvarez, J. & Schwartz, N. B. Measurement of Serum Steroid and Gonadotropin Levels and Uterine and Ovarian Variables throughout 4 Day and 5 Day Estrous Cycles in the Rat1. Biol. Reprod. 20, 659–670 (1979).

20. Smith, M. S., Freeman, M. E. & Neill, J. D. The control of progesterone secretion during the estrous cycle and early pseudopregnancy in the rat: Prolactin, gonadotropin and steroid levels associated with rescue of the corpus luteum of pseudopregnancy. Endocrinology 96, 219–226 (1975).

21. Butcher, R. L., Collins, W. E. & Fugo, N. W. Plasma concentration of LH, FSH, prolactin, progesterone and estradiol-17beta throughout the 4-day estrous cycle of the rat. Endocrinology 94, 1704–1708 (1974).

22. Heape, W. The ‘sexual season’ of mammals and the relation of the ‘pro-estrum’ to menstruation. Q. J. Microsc. Sci. 44, 1–70 (1900).

23. Stockard, C. R. & Papanicolaou, G. N. The existence of a typical oestrous cycle in the guinea pig with a study of its histological and physiological changes. Am. J. Anat. 22, 225–283 (1917).

24. Long, J. A. & Evans, H. M. The oestrous cycle in the rat and its associated phenomena. (University of California Press, 1922).

25. Everett, J. W. Progesterone and estrogen in the experimental control of ovulation time and other features of the estrous cycle in the rat. Endocrinology 43, 389–405 (1948).

26. Becker, J. B. et al. Strategies and methods for research on sex differences in brain and behavior. Endocrinology 146, 1650–1673 (2005).

27. Yener, T., Turkkani Tunc, A., Aslan, H., Aytan, H. & Caliskan, A. C. Determination of oestrous cycle of the rats by direct examination: how reliable? Anat. Histol. Embryol. 36, 75–77 (2007).

28. Marcondes, F. K., Bianchi, F. J. & Tanno, A. P. Determination of the estrous cycle phases of rats: some helpful considerations. Braz. J. Biol. 62, 609–614 (2002).

29. Paccola, C. C., Resende, C. G., Stumpp, T., Miraglia, S. M. & Cipriano, I. The rat estrous cycle revisited: a quantitative and qualitative analysis. Anim. Reprod., v vol. 10 (2018).

30. Hubscher, C. H., Brooks, D. L. & Johnson, J. R. A quantitative method for assessing stages of the rat estrous cycle. Biotech. Histochem. 80, 79–87 (2005).

31. Byers, S. L., Wiles, M. V., Dunn, S. L. & Taft, R. A. Mouse estrous cycle identification tool and images. PLoS One 7, 35538 (2012).

32. Wolcott, N. S. et al. Automated classification of estrous stage in rodents using deep learning. bioRxiv 2022.03.09.483678 (2022).

33. Sano, K. et al. Deep learning-based classification of the mouse estrous cycle stages. Sci. Rep. 10, (2020).

34. Barker, T. E. & Walker, B. E. Initiation of irreversible differentiation in vaginal epithelium. Anat. Rec. 154, 149–159 (1966).

35. Buchanan, D. L. et al. Role of stromal and epithelial estrogen receptors in vaginal epithelial proliferation, stratification, and cornification. Endocrinology 139, 4345–4352 (1998).

36. Galand, P., Leroy, F. & Chrétien, J. Effect of oestradiol on cell proliferation and histological changes in the uterus and vagina of mice. J. Endocrinol. 49, 243–252 (1971).

37. Montes, G. S. & Luque, E. H. Effects of ovarian steroids on vaginal smears in the rat. Cells Tissues Organs 133, 192–199 (1988).

38. Leroy, F., Galand, P. & Chrétien, J. The mitogenic action of ovarian hormones on the uterine and the vaginal epithelium during the oestrous cycle in the rat: a radioautographic study. J. Endocrinol. 45, 441–447 (1969).

39. Shorr, E. A New Technic for Staining Vaginal Smears: III, a Single Differential Stain Author (s): Ephraim Shorr Published by: American Association for the Advancement of Science Stable URL: https://www.jstor.org/stable/1667807. Science 94, p545–546 (1941).

40. Schank, J. C. Do Norway rats (Rattus norvegicus) synchronize their estrous cycles? Physiol. Behav. 72, 129–139 (2001).

41. Bartos, L. Vaginal impedance measurement used for mating in the rat. Lab. Anim. 11, 53–55 (1977).

42. Ramos, S. D., Lee, J. M. & Peuler, J. D. An inexpensive meter to measure differences in electrical resistance in the rat vagina during the ovarian cycle. J. Appl. Physiol. 91, 667–670 (2001).

43. Jaramillo, L. M., Balcazar, I. B. & Duran, C. Using vaginal wall impedance to determine estrous cycle phase in Lewis rats. Lab Anim. (NY). 41, 122–128 (2012).

44. Chesney, K. L., Chang, C. & Bryda, E. C. Using vaginal impedance measurement to identify proestrus in rats given luteinizing hormone releasing hormone (lhrh) agonist. J. Am. Assoc. Lab. Anim. Sci. 59, 282–287 (2020).

45. Singletary, S. J. et al. Lack of correlation of vaginal impedance measurements with hormone levels in the rat. Contemp. Top. Lab. Anim. Sci. 44, 37–42 (2005).

46. Cossio, R., Carreira, M. B., Vásquez, C. E. & Britton, G. B. Sex differences and estrous cycle effects on foreground contextual fear conditioning. Physiol. Behav. 163, 305–311 (2016).

47. Beatty, W. W. & Beatty, P. A. Hormonal determinants of sex differences in avoidance behavior and reactivity to electric shock in the rat. J. Comp. Physiol. Psychol. 73, 446–455 (1970).

48. Beatty, W. W. & Fessler, R. G. Gonadectomy and sensitivity to electric shock in the rat. Physiol. Behav. 19, 1–6 (1977).

49. Beatty, W. W. & Fessler, R. G. Sex differences in sensitivity to electric shock in rats and hamsters. Bull. Psychon. Soc. 10, 189–190 (1977).

50. Cooke, P. S., Spencer, T. E., Bartol, F. F. & Hayashi, K. Uterine glands: Development, function and experimental model systems. Mol. Hum. Reprod. 19, 547–558 (2013).

51. Woitowich, N. C., Beery, A. & Woodruff, T. A 10-year follow-up study of sex inclusion in the biological sciences. 1–8 (2020).

52. Brenner, R. M. & West, N. B. Hormonal regulation of the reproductive tract in female mammals. Annual review of physiology vol. 37 273–302 (1975).

53. Dixon, D. et al. Nonproliferative and proliferative lesions of the rat and mouse female reproductive system. J. Toxicol. Pathol. 27, (2014).

54. Sato, J., Nasu, M. & Tsuchitani, M. Comparative histopathology of the estrous or menstrual cycle in laboratory animals. J. Toxicol. Pathol. 29, 155–162 (2016).

55. Blume, S. R., Padival, M., Urban, J. H. & Rosenkranz, J. A. Disruptive effects of repeated stress on basolateral amygdala neurons and fear behavior across the estrous cycle in rats. Sci. Rep. 9, 1–18 (2019).

56. Blume, S. R. et al. Sex-And estrus-dependent differences in rat basolateral amygdala. J. Neurosci. 37, 10567–10586 (2017).

57. Gruene, T. M., Flick, K., Stefano, A., Shea, S. D. & Shansky, R. M. Sexually divergent expression of active and passive conditioned fear responses in rats. Elife 4, 1–9 (2015).

58. Iwata, J., Chida, K. & LeDoux, J. E. Cardiovascular responses elicited by stimulation of neurons in the central amygdaloid nucleus in awake but not anesthetized rats resemble conditioned emotional responses. Brain Res 418, 183–188 (1987).

59. Bakin, J. S., Lepan, B. & Weinberger, N. M. Sensitization induced receptive field plasticity in the auditory cortex is independent of CS-modality. Brain Res. 577, 226–235 (1992).

60. Erickson, M. T. & Walter, E. T. Differential expression of pseudoconditioning and sensitization by siphon responses in Aplysia: Novel response selection after training. J. Neurosci. 8, 3000–3010 (1988).

61. Zambetti, P. R. et al. Pavlovian fear conditioning does not readily occur in rats in naturalistic environments. bioRxiv 2021.10.20.465116 (2021) doi:10.1101/2021.10.20.465116.

62. Frye, C. A., Petralia, S. M. & Rhodes, M. E. Estrous cycle and sex differences in performance on anxiety tasks coincide with increases in hippocampal progesterone and 3α,5α-THP. Pharmacol. Biochem. Behav. 67, 587–596 (2000).

63. Mora, S., Dussaubat, N. & Díaz-Véliz, G. Effects of the estrous cycle and ovarian hormones on behavioral indices of anxiety in female rats. Psychoneuroendocrinology 21, 609–620 (1996).

64. Bitran, D., Purdy, R. H. & Kellog, C. K. Anxiolytic effect of progesterone is associated with increases in cortical alloprenanolone and GABAA receptor function. Pharmacol. Biochem. Behav. 45, 423–428 (1993).

65. Marcondes, F. K., Miguel, K. J., Melo, L. L. & Spadari-Bratfisch, R. C. Estrous cycle influences the response of female rats in the elevated plus-maze test. Physiol. Behav. 74, 435–40 (2001).

66. Walf, A. A. & Frye, C. A. Antianxiety and antidepressive behavior produced by physiological estradiol regimen may be modulated by hypothalamic-pituitary-adrenal axis activity. Neuropsychopharmacology 30, 1288–1301 (2005).

67. McClintock, M. & LeFevre, J. Reproductive senescence in female rats: a longitudinal study of individual differences in estrous cycles and behavior. Biol. Reprod. 38, 780–789 (1988).

68. Lu, K. H., Hopper, B. R., Vargo, T. M. & Yen, S. S. C. Chronological Changes in Sex Steroid, Gonadotropin and Prolactin Secretion in Aging Female Rats Displaying Different Reproductive States1. Biol. Reprod. 21, 193–203 (1979).

69. Yin, W. et al. Testing the Critical Window Hypothesis of Timing and Duration of Estradiol Treatment on Hypothalamic Gene Networks in Reproductively Mature and Aging Female Rats. 1–16 (2015) doi:10.1210/en.2015-1032.

70. Fiala, J. C. Reconstruct: a free editor for serial section microscopy. J Microsc 218, 52–61 (2005).

71. Ostroff, L. E., Cain, C. K., Bedont, J., Monfils, M. H. & LeDoux, J. E. Fear and safety learning differentially affect synapse size and dendritic translation in the lateral amygdala. Proc. Natl. Acad. Sci. U. S. A. 107, 9418–9423 (2010).

